# Phylogeny of the species-rich *Pilea* Lindl. (Urticaceae) supports its revised delimitation and infrageneric classification, including the resurrection of *Achudemia* Blume

**DOI:** 10.1101/2020.07.01.180109

**Authors:** Long-Fei Fu, Fang Wen, Olivier Maurin, Michele Rodda, Yi-Gang Wei, Alexandre K. Monro

## Abstract

*Pilea* Lindl., with 933 published names is the largest genus within the Urticaceae. *Pilea* was last monographed in 1869 and whilst the monophyly of the genus has been proposed by previous authors, this has been based on incomplete taxon sampling and the failure to resolve the position of key taxa. We aimed to generate a robust phylogeny for *Pilea* and allied genera that could provide a framework for testing the monophyly of *Pilea*, revising its delimitation and for answering broader scientific questions about this species-rich genus. To do so, we sought to sample taxa representative of previous infrageneric classifications and with anomalous inflorescences or flower configurations and to use the resulting phylogeny to evaluate the delimitation of *Pilea* and to establish an infrageneric classification. In addition, we included a representative of the Polynesian genus *Haroldiella* which, morphologically, is very similar to *Pilea*. Using Sanger sequence data from two plastid and one nuclear regions we constructed a phylogeny using Bayesian Inference, Maximum Likelihood and Maximum Parsiomony approaches. We used our phylogeny to evaluate the informativness of 19 morphological traits and applied both to delimit a monophyletic genus and infrageneric sections. Our results recovered *Pilea* as paraphyletic with respect to *Lecanthus*, a consequence of the recovery of a monophyletic clade comprising sections *Achudemia* and *Smithiella*, neither of which had been adequately sampled in previous studies. We also recovered *Pilea* as polyphyletic with respect to *Haroldiella*. We identified isomery between male and female flowers, flower part number and male sepal arrangement as being phylogenetically informative traits that can be used to delimit two genera, *Achudemia*, including section *Smithiella*, recovered as sister to *Lecanthus*, and *Pilea*, including *Haroldiella*, recovered as sister to both. On the basis of our evaluation of both morphological traits and phylogenetic relationships we propose a new infrageneric classification for the genus comprising seven sections, five of which we describe for the first time, § *Trimeris* Y.G.Wei & A.K.Monro, § *Lecanthoides* C.J.Chen, § *Angulata* L.F.Fu & Y.G.Wei, § *Tetrameris* C.J.Chen, § *Verrucosa* L.F.Fu & Y.G.Wei, § *Plataniflora* L.F.Fu & Y.G.Wei and § *Leiocarpa* L.F.Fu & Y.G.Wei. We also identify a trend of decreasing merism and fruit size, and increasing species-richness as *Pilea* diverges. In addition, we recover strong geographical structure within our phylogeny, sufficient to propose that *Pilea* originated in the IndoMalaya biogeographic domain.

## 1. Introduction

*Pilea* Lindl., with 933 published names (IPNI, 2020), 604 accepted names (WCVP, 2020) and likely 715 species worldwide (Monro, 2004) is the largest genus within the Urticaceae and has a pantropical and subtropical distribution. *Pilea* is characterized by succulent herbs, shrubs and epiphytes whose flowers are wind pollinated, opening explosively, and seed which is mechanically dispersed over short distances through the reflexing of the staminodes. It is most species-rich in forested rocky habitats, especially on limestone or ultramafic rocks at elevations between of 500 and 2,000 masl, in the Greater Antilles, Central America and the Andes. Members of the genus may be distinguished from other genera in the family by the combination of opposite (rarely alternate) leaves, intrapetiolar stipules, an absence of stinging hairs, male inflorescences not fused to form a receptacle-like structure, and free female sepals (Fig. 1 & Fig. 2). As is the case for many species-rich genera, *Pilea* has not been monographed since the 19thC (Weddell, 1869) at which time the genus comprised ca 150 spp‥ Instead, its taxonomy has been revised piecemeal through flora treatments (Monro, 2006).

**Fig. 1.**
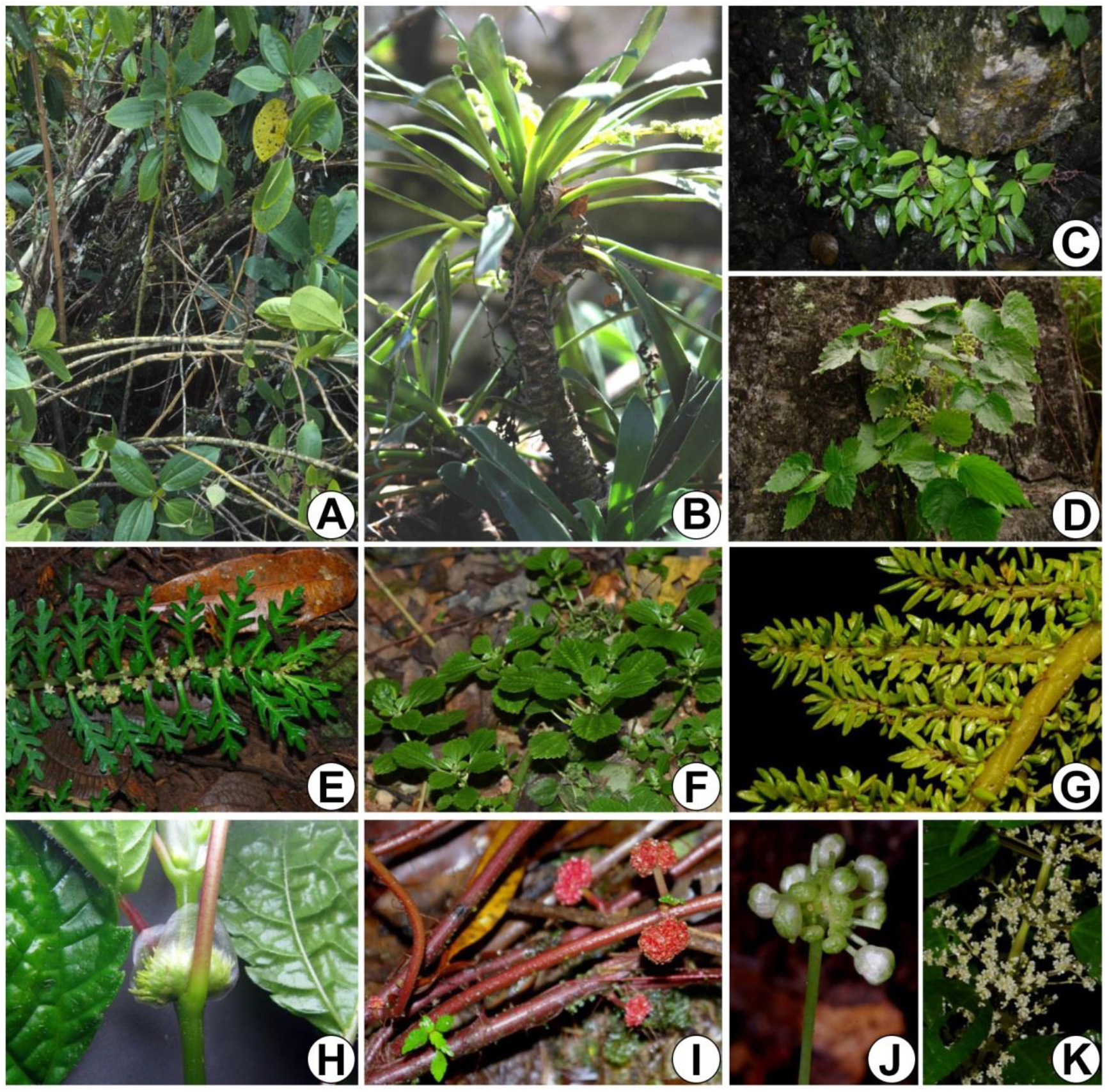
Morphological diversity of *Pilea* and *Achudemia*. A, *P. longicaulis* (shrub); B, *P. fairchildiana* (shrub, with alternate, spirally arranged leaves); C, *A. boniana* (herb, epipetric); D, *P. paniculigera* (herb, epipetric); E, *P. matama* (unequal opposite leaves, epiphytic with capitate female inflorescences); F, *P. peploides* (herb, clumped); G. *P. sp* aff. *microphylla*; H, *P. rivularis* (female inflorescence enclosed by stipules); I, *P. aff. pittieri* (herb, male capitate inflorescence arising from stolons); J, *P. angustifolia* (herb, male capitate inflorescence); *P. notata* (male cyme inflorescence). A-B, E, G-J were photographed by Alexandre K. Monro; D was photographed by Yi-Gang Wei; C-D, F, K were photographed by Long-Fei Fu.

**Fig. 2.**
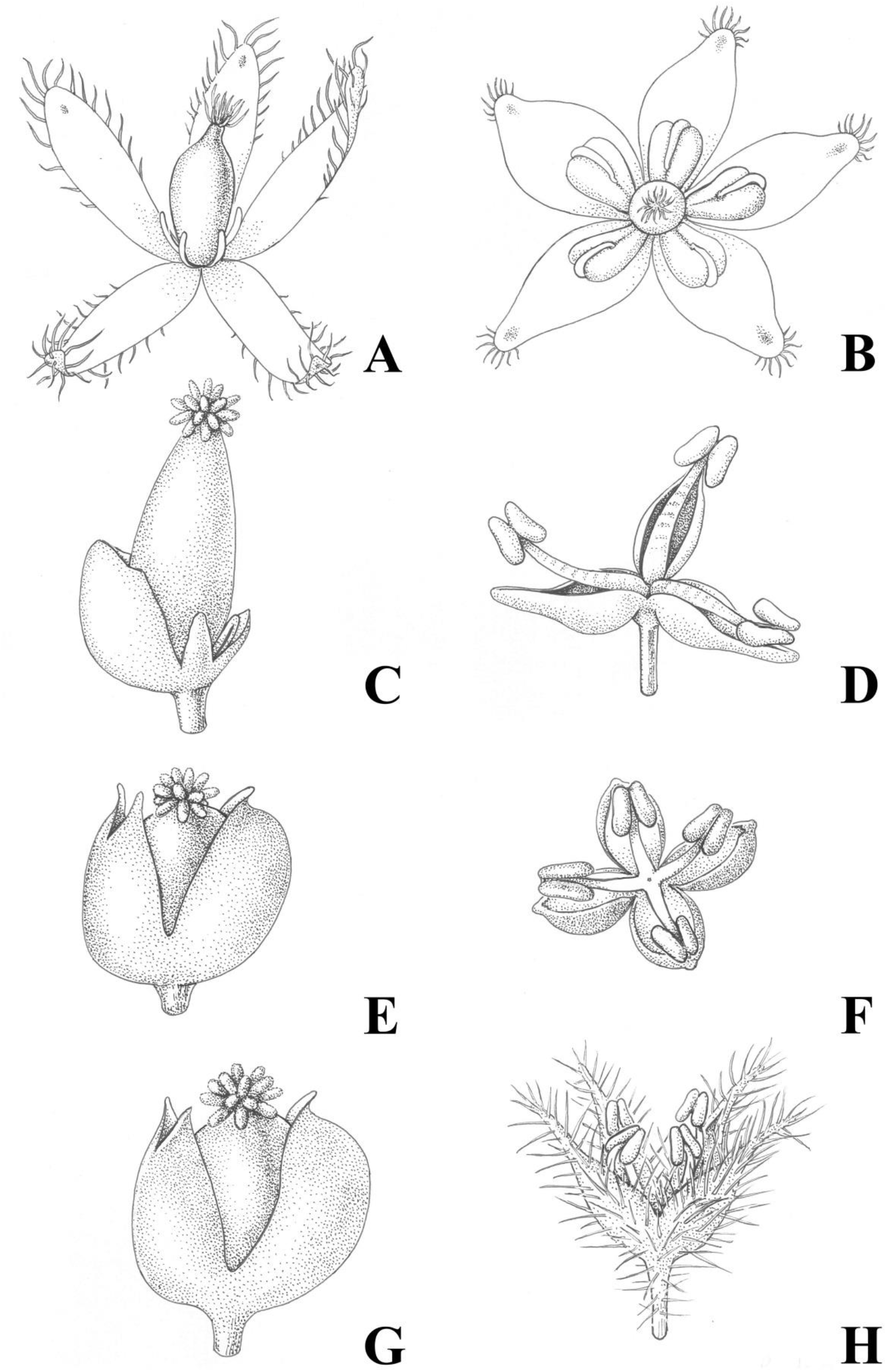
Illustration of male and female flowers of *Pilea* and *Achudemia*. A, *A. javanica* (female flower); B, *A. javanica* (male flower); C, *P. tripartite* (female flower); D, *P. tripartite* (male flower); E, *P. plataniflora* (female flower); F, *P. plataniflora* (male flower); G, *P. microphylla* (female flower); H, *P. microphylla* (male flower). Illustration by Margaret Tebbs.

*Pilea* belongs to the Elatostemaea tribe (Gaudichaud, 1830) which, including *Sarcopilea* Urb., has been recovered as monophyletic and sister to *Lecanthus* Wedd. (Monro, 2006; Jestrow et al., 2012; Wu et al., 2013; Tseng et al. 2019). The Elatostemeae comprises mainly succulent, shade-loving, wind-pollinated species which, as is the case for each tribe in the family, show a great variation in female inflorescence arrangement, ranging from open panicles to spikes and fused receptacle-like structures. Despite several molecular studies, doubts remain over the status of *Achudemia* Blume. Different accessions of *A. japonica*, having been recovered within, or sister to, *Pilea* (Monro, 2006) and currently it is included within *Pilea* (Friis, 1989; Chen and Monro, 2003). In addition, neither the Polynesian endemic, *Haroldiella* J.Florence, whose morphological circumscription is congruent with *Pilea*, or section *Smithiella*, characterised by strongly asymmetrical spicate inflorescences, were sampled in previous studies, suggesting that the monophyly of the genus remains untested.

*Achudemia*, currently treated as a section of *Pilea,* comprises four species of herb that grow in deep forest shade, stream sides, gorges and caves in Indomalaya. *Achudemia* was established by Blume (1856: 57) to account for a *Pilea*-like collection from Java (Indonesia) which had bisexual (hermaphrodite) five-parted flowers. It appears that Blume described the flowers as bisexual in error as neither the holotype, type illustration, or any other collections have been observed to have bisexual flowers.

*Pilea* section *Smithiella*, comprises a single species of herb from Indomalaya, also growing in deep shade (Chen, 1995; Chen and Monro, 2003). *Smithiella* was generated by Dunn (1920) to account for *Pilea*-like material from the Eastern Himalayas characterised by strongly asymmetrical spicate inflorescences of five-parted flowers. Dunn had been unaware of an earlier homonym with priority, *Smithiella* H. Perag. & Perag. and in 1981, Bennet (1981) created a replacement name, *Aboriella*.

*Haroldiella* comprises two species from Austral Polynesia growing on rocky outcrops or in rain forest*. Haroldiella* was described by Florence (1997) based on plants with alternate, spirally arranged pinnately nerved leaves. As with all *Pilea* from French Polynesia (Florence, 1997), they also share the trait of two-sepalate female flowers, a condition very rare elsewhere in the genus. With the recovery of *Sarcopilea domingensis* Urb., a taxon with spirally arranged, pinnately nerved alternate leaves, within a monophyletic *Pilea* (Monro, 2006, Jestrow et al., 2012), the characters used to delimit *Haroldiella* no longer support its separation as a distinct genus.

Both *Achudemia* and *Aboriella* differ from other *Pilea* species in having male and female flowers with five sepals (Fig. 2). Where free, the same number of perianth parts in male and female flowers is uncommon within the Urticaceae and within *Pilea*, it is a condition restricted to a basal, relatively species-poor clade (Monro, 2006) comprising Afrotropical, Indomalayan and neotropical species.

Previous phylogenetic studies have consistently recovered *Lecanthus* as the genus most closely related to *Pilea* (Monro, 2006; Wu et al., 2013, 2018), from which it be distinguished by its male inflorescences being fused to form a concave receptacle-like structure reminiscent of *Elatostema* J.R.Forst. & G.Forst. It also has an equal number of male and female perianth parts, either four or five (Chen and Monro, 2003).

Given the above, and with the limited sampling of *Pilea* species with anomalous inflorescence arrangements (spicate, receptacle-like) or flower-part number (five), together with the ambiguous position of *Achudemia*, generic delimitation is potentially unstable. Furthermore, the two main infrageneric classifications of *Pilea* (Weddell, 1856; Chen, 1982) have been demonstrated to be largely para- or polyphyletic.

For the above reasons, we aimed to generate a robust phylogeny for *Pilea* and allied genera that could provide a framework for revising the delimitation of the genus and the answering of broader scientific questions about this species-rich and poorly studied genus. To do so we sought to increase taxon sampling for the genus, encompassing all previous sections, and the full range of morphological variation and geographical occurrence, using an evaluation of the informativeness of morphological traits support the establishment of an infrageneric classification.

## 2. Materials and methods

### 2.1 Taxon sampling

We included 137 accessions representing 125 taxa (Table 1). These included 18 outgroup taxa from the Cannabaceae, Moraceae and representatives of all Urticaceae tribes except for Cecropieae (see Table 1) and 107 ingroup taxa (*Lecanthus* + *Pilea*). Within the Elatostemeae, all genera were sampled except for the monotypic *Metapilea* which is likely extinct (Wu et al., 2018), known only from the type and could not be sampled. This encompassed the following taxa, *Elatostema* (3 spp), *Elatostematoides* (1 sp), *Lecanthus* (2 spp), *Pilea* (105 spp), *Procris* (1 sp) and *Polychroa* (1 sp). We focussed on the ITS nuclear region and the *rbcL* and *trnL-F* plastid regions. We combined sequences generated by previous studies (Monro, 2006; Jestrow et al., 2012; Wu et al., 2013, 2018; Kim et al., 2015) 141 sequences of which were generated by ourselves, excluding those accessions where we felt that the identifications were ambiguous, or where sequence data for only a single region could be obtained. An exception was made for the single sequence of *Haroldiella* that we were able to obtain. Our sampling of *Pilea* included representatives of all infrageneric sections proposed by Chen (1982, Chen and Monro, 2003). We did not structure our sampling to include representatives of Weddell’s sections as these were all demonstrated to be para- and polyphyletic by Monro (2006).

**Table 1.** Statistics for the molecular datasets used in this study.

|  | Number of<br>sequences<br>(ingroup/outgroup) | Aligned<br>length<br>(bp) | Length<br>variation<br>(bp) | Variable<br>characters<br>(bp) | Parsimony-infor<br>mative characters<br>(bp) | Model<br>selected<br>(AIC) |
| --- | --- | --- | --- | --- | --- | --- |
| ITS | 119/18 | 716 | 472-588 | 490 | 398 | GTR+I+G |
| *trnL-trnF* | 117/18 | 1059 | 402-1059 | 532 | 356 | GTR+I+G |
| *rbcL* | 61/18 | 637 | 629-637 | 137 | 82 | GTR+I+G |
| Combined plastid | 119/18 | 1696 | 1038-1696 | 669 | 438 | GTR+I+G |
| Combined all | 119/18 | 2412 | 1572-2231 | 1159 | 836 | GTR+I+G |

Sequence data were obtained for the four species of *P.* sect. *Achudemia*, the monotypic *P.* sect. *Smithiella*, ten from *P.* sect. *Tetrameris* (approximately 2/3 of the species), 85 from *P.* sect. *Urticella* (approximately ¼ of the species), seven from *P.* sect. *Pilea* (approximately 2/3 of the species), two from *P.* sect. *Dimeris* (1/2 of the species) and one from *P.* sect. *Lecanthoides* (1/2 of the species). Four species of Moraceae (*Fatoua villosa* Nakai, *Morus alba* L., *Sorocea affinis* Hemsl., *Trophis racemosa* (L.) Urb.) and two species of Cannabaceae (*Cannabis sabiva* L., *Humulus lupulus* L.) were chosen as outgroups based on the previous analyses (Zhang et al., 2011; Kim et al., 2015). Species names, the accession numbers of sequences downloaded from GenBank, and newly generated sequences used in this study are listed in Supplementary Text 1.

### 2.2 DNA isolation, PCR amplification and sequencing

Genomic DNA was extracted from fresh or dried materials using a modified CTAB protocol (Chen et al., 2014). The nrITS region was amplified using primers ITS 4 and ITS 5 (White et al., 1990) and *rbcL* using primers 1F and 724R (Fay et al., 1997). The *trnL-F* spacer was amplified using primers e and f (Taberlet et al., 1991) for most accessions while for few problematic cases, we employed primers c and d to separate *trnL-F* into two overlap regions then concatenated sequences (Taberlet et al., 1991). The PCR amplification were set at 94 °C for 5 min, 30 cycles of 94 °C for 30 s, 55 °C for 30 s, and 72 °C for 45 s and a final extension at 72 °C for 10 min. The PCR products were checked on 1% agarose gels before being purified using a QiaQuick gel extraction kit (Qiagen, Inc., Valencia, California, USA) and directly sequenced in both directions using the amplification primers on an ABI 3730 automated sequencer (Applied Biosystems, Forster City, California, U.S.A.).

### 2.3 Phylogenetic analyses

Raw sequences were edited and assembled using the software Lasergene Navigator (DNAStar, Madison, Wisconsin, USA) with subsequent manual adjustments. The output DNA sequences were then aligned using MAFFT version 7.0 (Katoh and Standley, 2013) with default settings, followed by manual adjustment. The three datasets (nrITS, *rbcL*, and *trnL-F* spacer) were aligned independently. Alignments were adjusted manually in MEGA 5.1 (Tamura et al., 2011). Phylogenies were reconstructed based on the nrITS dataset, the combined plastid datasets (*rbcL*, and *trnL-F* spacer), and all three datasets combined (nrITS, *rbcL*, and *trnL-F* spacer), respectively. All of these reconstructions were analysed using Bayesian inference (BI), maximum likelihood (ML), and maximum parsimony (MP) methods. A visual comparison of the two best tree topologies generated by ML analyses of cpDNA and nrITS datasets were performed to compare topological incongruence. A conflict in tree topologies of each tree was considered significant when incongruent topologies both received bootstrap values ≥ 80% (Monro, 2006; Tseng et al., 2019).

Best-fit DNA substitution models were selected using the Akaike Information Criterion (AIC) in Modeltest v 2.7 (Posada and Crandall, 1998) for each data partition. The substitution model of the sequences was set to GTR+G+I for each single dataset based on Modeltest. BI analyses were based on a Markov chain algorithm implemented in MRBAYES 3.2.6 (Huelsenbeck and Ronquist, 2001).

ML analyses with 1000 bootstrap resampling (MLBS) were conducted using the online version of RAxML-HPC2 v8.2.9 (Stamatakis et al., 2008) available at the CIPRES Science Gateway version 3.3 (http://www.phylo.org/index.php/portal/) (Miller et al., 2010) with the gamma model of rate heterogeneity.

MP analyses were performed using PAUP* v4.0b10 (Swofford, 2002), in which all characters were unordered and equally weighted, and gaps were treated as missing data. Heuristic searches of MP were conducted with 100 random addition replicates with tree tree-bisection–reconnection (TBR) branch swapping and MulTrees in effect. Branch supports were assessed using 1000 bootstrap replicates (maximum parsimony bootstrap; MPBS) with the sample settings the same as those for heuristic searches.

### 2.4 Estimates of support

In Bayesian analyses, posterior probabilities (PP) below 0.9 were considered as providing no support, between 0.9 and 0.94 as providing weak support, between 0.95 and 0.99 as providing moderate support, and 1.0 as providing strong support (Tseng et al., 2019).

In bootstrap analyses of the ML (BSML) and MP (BSMP) analyses, values below 70% were considered as providing no support, between 70-79% as providing weak support, between 80-89% as providing moderate support, and 90-100% as providing strong support (Tseng et al., 2019).

### 2.5 Morphological trait evolution

Based on existing phylogenetic studies we performed ancestral state reconstructions (ASR) in order to evaluate the phylogenetic informativness of selected morphological traits and so apply these to the delimitation of *Pilea* and allied genera, and the establishment of an infrageneric classification of *Pilea*. Our aim was to establish a classification that was both phylogenetically congruent and morphologically diagnosable.

Nineteen morphological traits were coded for analysis (see Supplementary Text 2). Traits were selected on the basis that they had been used in previous classifications and revisions of *Pilea* (Weddell, 1856; Chen, 1982; Monro, 2006, 2015). Traits were scored based on the examination of herbarium specimens and description in the literatures (Chen, 1982; Friis, 1989; Monro, 1999, 2015; Chen and Monro, 2003; Monro et al., 2012; Fu et al., 2017a; Yang et al., 2018).

Likely transitions between trait states through evolution were reconstructed using ML methods in Mesquite v.3.51 (Maddison and Maddison, 2015). We sampled the last 1000 trees from the post burn-in set of the Bayesian analysis using combined dataset and an equal rate model (Mk1) was selected for all traits. To account for phylogenetic uncertainty, we used ‘Trace character over trees’. All reconstructions were integrated over the 1000 trees from the post burn-in set and summarized on one of these trees that most matched our hypothesized topology. The results were summarized as a percentage of changes of trait states using the option of ‘Average frequencies across trees’.

### 2.6 Delimitation of infrageneric groupings

Given the number of species in *Pilea*, an infrageneric classification can be a practical way to ease identification, as well as providing a framework for answering broader evolutionary questions. With these aims our classification needed to reflect both phylogenetic relationships and be morphologically diagnosable. We decided to base our classification of *Pilea* on sections rather than subgenera as the distinction between the two is unclear (Brizicky, 1969) and in this way we maintain the terminology adopted by Weddell (1856, denoted by the symbol ‘§’) and Chen (1982). We have also aimed to establish sections in accordance with the International Code of Nomenclature for algae, fungi and plants (Turland et al., 2018).

## 3. Results

### 3.1. Phylogenetic reconstruction

Characteristics and statistics of the datasets used in this study are summarized in Table 1. The comparison of trees for cpDNA (*trnL-F*, *rbcL*) and nrITS revealed an incongruence between the outgroup taxa (Figs. S1 & S2). Because this incongruence did not affect the topology of the ingroup taxa and the phylogeny of the combined dataset showed better resolved trees with higher support values, than individual trees, we used the combined dataset for subsequent analyses, including that of transitions between morphological trait states. The ingroup taxa were recovered as monophyletic (Fig. 3) with strong support (PP1.0/BSML100%/BSMP100%).

**Fig. 3.**
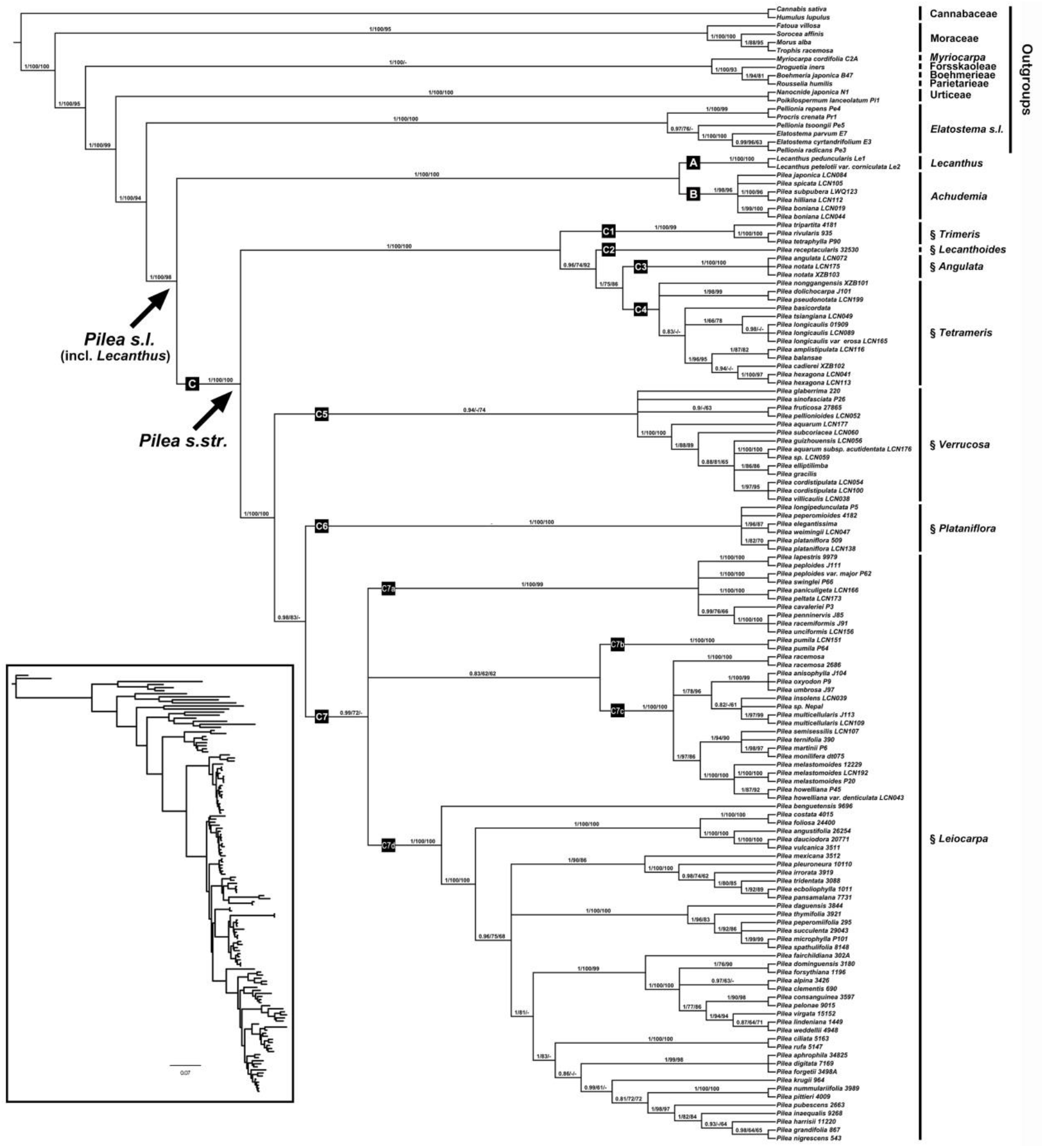
Phylogenetic tree of *Pilea* generated from Bayesian Inference (BI) of combined dataset (nrITS, *trnL-F* spacer and *rbcL*). Numbers on the branches indicate the posterior probability (≥0.8) of BI and bootstrap values (≥60%) of the maximum likelihood (ML) and the maximum parsimony (MP) analyses.

### 3.2. Phylogenetic relationships of Pilea

*Pilea*, including *Achudemia* and *Haroldiella* was recovered as paraphyletic with respect to *Lecanthus* (Fig. 3 & Fig. S3). Two strongly supported clades attributable to *Pilea sensu latu* were recovered. The first (Fig. 3, Clade B, labelled as *Achudemia*), was recovered sister to *Lecanthus* (Clade A) and included all accessions from *P.* sect. *Achudemia* and *P.* sect. *Smithiella* (*P. subpubera*, *P. boniana*, *P. hilliana*, *P. japonica*, *P. spicata*) with strong support (PP1.0/BSML100%/BSMP100%). The second clade (Fig. 3, Clade C) comprised all other accessions of *Pilea* with strong support (PP1.0/BSML100%/BSMP100%) and, in the analysis of the ITS sequence data (Fig. S3), *Haroldiella*, with strong support (1/100/100). Within Clade C, seven subclades were recovered with strong to weak support (C1 (1/100/99), C2 (0.96/74/92), C3 (1/100/100), C4 (-/-/-/), C5 (0.94/-/74), C6 (1/100/100) and C7 (0.99/72/-)) (Fig. 3). *Haroldiella* was recovered within clade C7d (Fig. S3, 1/100/98). Clade C4 and C5 were no and weakly supported by all methods used to analyse the data, albeit they comprise groups united by the morphological trait states of four-parted female flower and ornamented achenes, respectively. Clade C7 was recovered with strong to weak support but comprised four strongly supported subclades (C7a (1/100/99), C7b (1/100/100), C7c (1/100/100), C7d (1/100/100)).

### 3.3. Geographical structure

Clade A (Figs. 4-5 & Fig. S23) comprises taxa with an Asia distribution, although *Lecanthus* also includes species (not sampled) from Africa. Clade B (Figs. 4-5 & Fig. S23) comprises taxa from East and Southeast Asia.

**Fig. 4.**
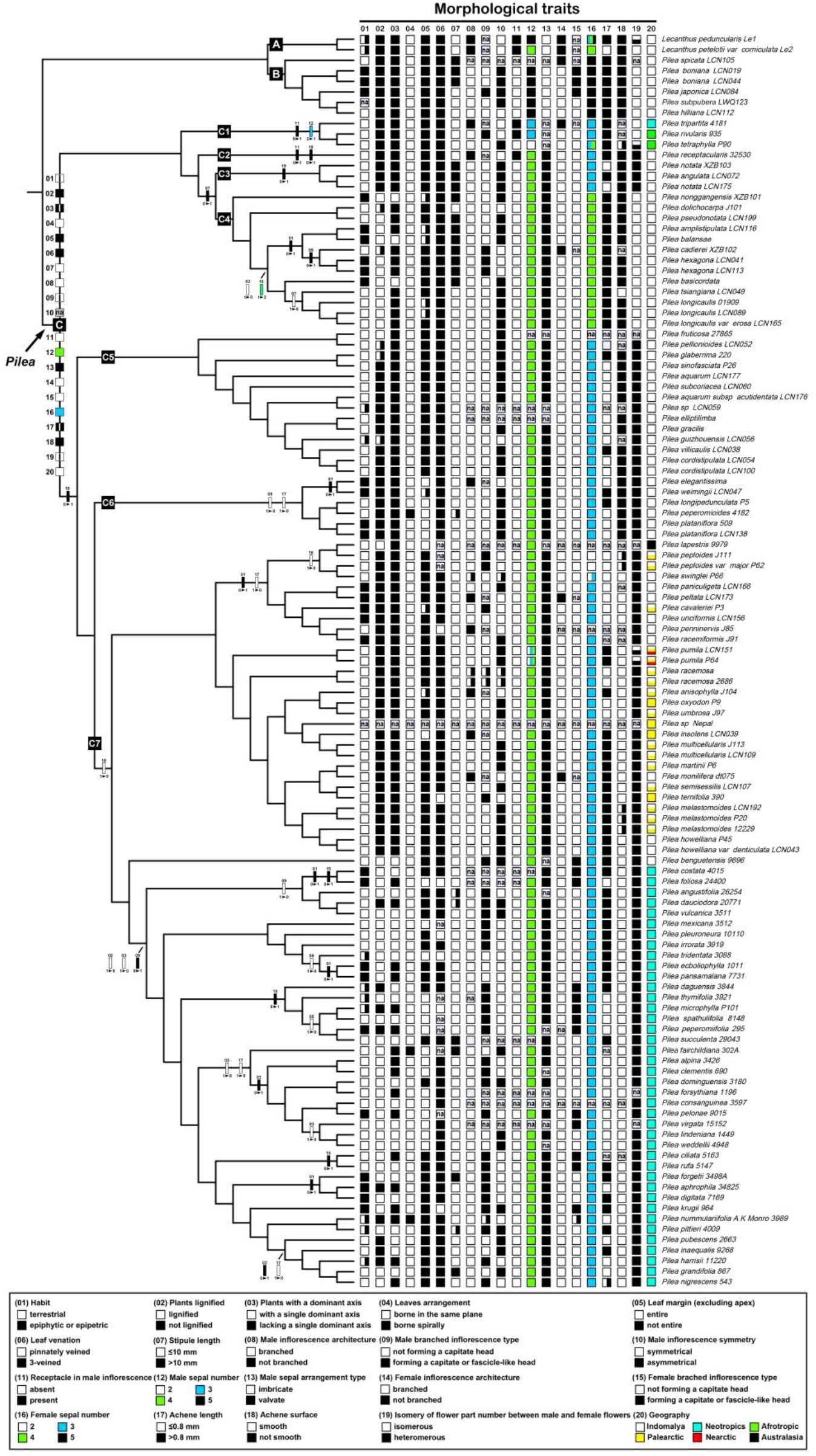
Reconstruction of the evolution of twenty morphological traits in *Pilea* based on our Bayesian Inference analysis of the combined dataset. The trait states at the *Pilea* node indicate the ancestral states of the genus. Transitions are indicated as filled boxes on the branches. Traits are shown above boxes and state transitions below. Descriptions of traits and their states are provided in the legend. The three clades (A–C) and seven subclades (C1-C7) correspond to those in Fig. 2. Reconstructions for each trait can be seen in Supplementary Figures S4 to S23.

**Fig. 5.**
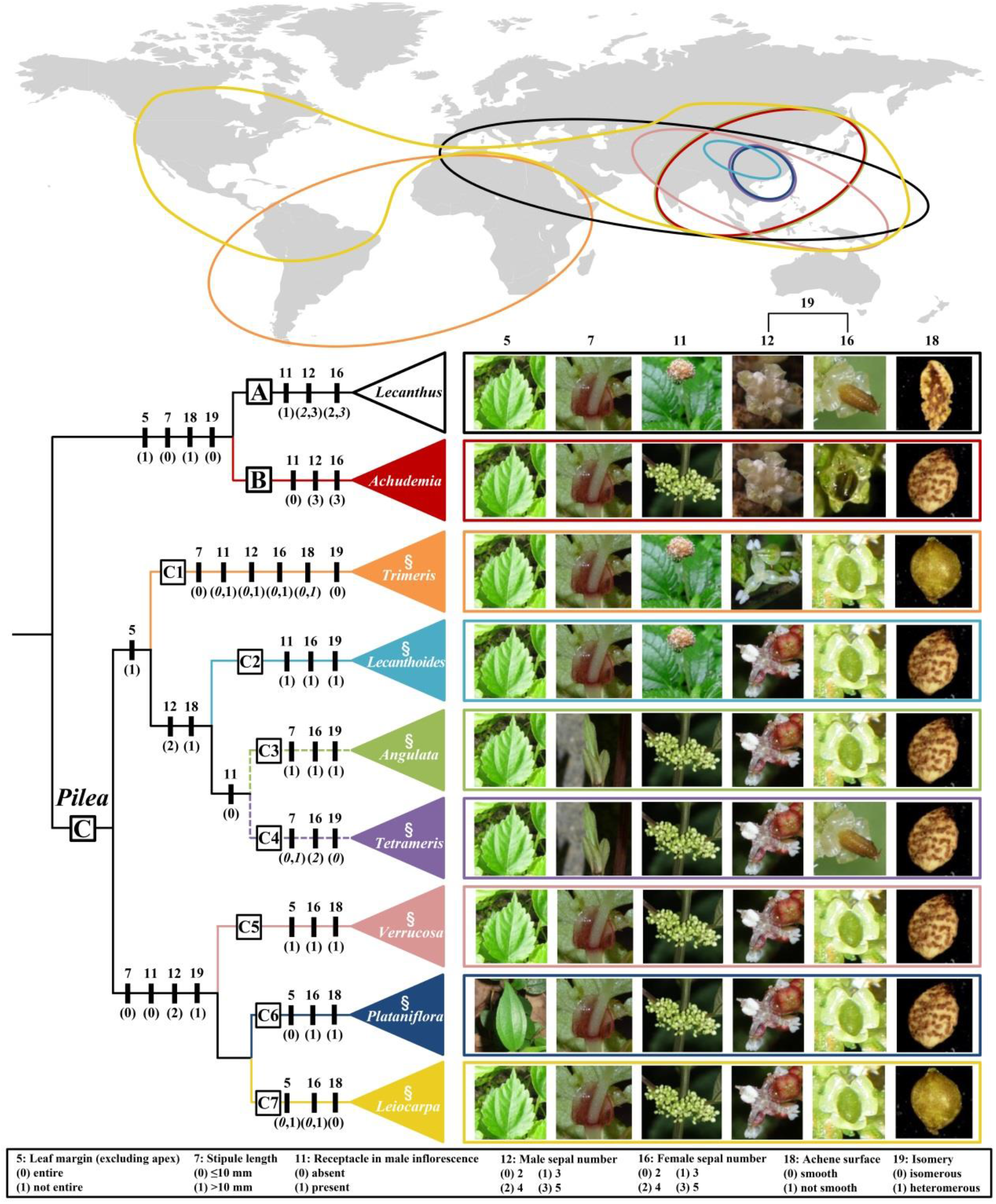
Proposed infrageneric classification of *Pilea* and their geographical distribution. Filled boxes illustrate all states for each section. *Derived* states in italics, ancestral in regular font (based on ancestral state reconstructions summarised in Fig. 3). Images illustrate the ancestral states (replaced by common state when ancestral one is not available) for each infrageneric section.

Clade C (Figs. 4-5 & Fig. S23) comprises species with a pantropical distribution except for Australia and New Zealand. Within clade C, subclade C1 comprises taxa from Africa, Asia and Latin America; subclades C2, C3, C4, C5 and C6 comprise taxa from East Asia; and subclade C7 taxa from the pantropics. *Haroldiella*, restricted to Polynesia was recovered within a polytomy within clade C7d meaning that it is more closely related to neotropical species, than to African or Asian ones. Within clade C7, three subclades show a strong geographical association. Subclade C7d, which harbours the greatest number of species, is strongly associated with the neotropics, the Greater Antilles, Andes and Central America in particular. Subclades C7b and C7c comprise predominantly palearctic taxa.

### 3.4. Morphological trait evolution

The 19 morphological traits were mapped onto the Bayesian Inference (BI) tree based on the Maximum Likelihood analyses (Fig. 4 & Figs. S4-S22). This recovered five-sepalate male and female flowers, and imbricate male flower sepals as synapomorphies for Clade A + B (*Achudemia*), with a reversal in *L. petelotii* var. *corniculata* which has four-sepalate male and female flowers. Achene bearing a crescent-shaped protuberance and unbranched male inflorescences were recovered as autoapomorphies for Clade A (*Lecanthus*). Five-sepalate female and male flowers were recovered as a plesiomorphy, and branched male inflorescences as a synapomorphy, for Clade B (*Pilea* section *Achudemia* + *P.* section *Smithiella*). The presence or absence of a crescent-shaped protuberance on the achene and branching, or not, of the male inflorescence enable clades A and B to be readily distinguished from each other.

Four plesiomorphies were recovered for Clade C (all remaining *Pilea* species), stipules ≤ 10 mm, male flowers 3-sepalate, achene > 0.8 mm, achene ornamented. These were manifested in the basal subclades, transforming to other states through the tree. All other traits were recovered as homoplastic for the clade.

Within subclades of Clade C, however, several plesiomorphic and synapomorphic trait states were recovered. For clade C1, we recovered male flowers 3-sepalate (with a reduction to two-sepalate for *Pilea tetraphylla*) as a synapomorphy and male sepals valvate as a plesiomorphy. For C2, we recovered branches of the male inflorescence fused to form a receptacle-like structure as a synapomorphy. For C3, we recovered stipules > 10 mm as a synapomorphy. For C5 we recovered achene surface ornamented as a plesiomorphy. For C6 we recovered achene surface ornamented and leaf margin entire as plesiomorphies. For C7 we recovered achene surface smooth as a plesiomorphy. For C4 we recovered no synapomorphies or plesiomorphies. It could, however, be morphologically diagnosed based on unique combinations of morphological traits (male and female flowers 4-sepalate), as could several other monophyletic groupings within clade C.

## 4. Discussion

### 4.1. Resurrection and expansion of Achudemia

Previous studies (Monro, 2006; Jestrow et al., 2012; Wu et al., 2013, 2015, 2018) did not attempt to establish formal infrageneric classifications because of limited morphological trait and taxon sampling, especially of the more basal taxa. Monro (2006) did, however, suggest a preliminary and informal classification of *Pilea* into six ‘units’ based on geography and isomery of flower part number between male and female flowers, cystolith distribution on the leaves and the presence of multicellular hairs. In this study we sought to propose a formal classification based increased taxon, morphological and geographical sampling. Our approach resolved the positions of all members of section *Achudemia*, including *Pilea subpubera*, synonym of *Achudemia javanica*, genus type for *Achudemia*, and *Pilea* (*Achudemia*) *japonica*, ambiguously recovered within and outside of *Pilea* by Monro (2006). In doing so recovered a paraphyletic *Pilea*. The paraphyly can be resolved through the exclusion of 5-sepalate male flowered taxa, formally assigned to Chen’s (Chen, 1982; Chen and Monro, 2003) sections *Achudemia* and *Smithiella*, into a resurrected and expanded genus, *Achudemia*, which can be distinguished from other *Pilea* species and *Lecanthus*, based on flower, male inflorescence and achene morphology.

*Haroldiella* was recovered within *Pilea*. This result was not surprising given that the justification for treating the taxa within a genus distinct to *Pilea* (Florence, 1997) was based on the presence of alternate, spirally arranged leaves, a trait state expressed elsewhere within *Pilea* (e.g. *P. peperomiodes*, *P. fairchildiana*).

Whilst almost all of the morphological trait states assessed were found to be homoplastic, our expanded sampling did enable us to recover seven morphologically diagnosable clades within *Pilea* and to use these to establish seven sections, partly congruent with those proposed by Chen (1982) on morphology alone.

### 4.2. Pilea originated in IndoMalaya

Mapped onto our phylogeny, geographical occurrence suggests IndoMalaya as the centre of origin for *Pilea* (Fig. 4 & Fig. S23)*, Lecanthus*, *Achudemia* and the basal clades of *Pilea* (C2 to C6) predominantly comprising Indomalayan species. This confirms the findings of Monro (2006) and Wu et al. (2018). Given the relationships between geographical areas suggested by our results, the most plausible scenario for the dispersal of *Pilea* is of two independent events. The first, early in the divergence of the genus (Fig. 4 & Fig. S23, clade C1) from IndoMalaya to Africa and the northern Neotropics which has resulted in a lineage with a modest number of species. The second, later in the divergence of the genus, involving dispersal to the Palearctic (Fig. 4 & Fig. S23, clade C7) and later from the Palearctic to the Neotropics (clade C7) resulting in species radiations in the Andes and Greater Antilles. According to the dated Urticaceae phylogeny of Wu et al. (2018, Fig. 1), this first dispersal event would have occurred ca 35 MYA (late Eocene) and the second ca 25 MYA (late Oligocene). A plausible mechanism and route for second dispersal to the Neotropics would be the Bering land bridge at some point between the late Cretaceous and late Neogene (Wen et al., 2016). Whilst long distance dispersal, invoked by Wu et al. (2018) may have played a role in the first.

### 4.3. Diversification accompanied by reductions in merism, achene size and ornamentation and an increase in species number

Chen (1982) proposed a reduction in female flower part number as an evolutionary trend in *Pilea* and our results support this, five-parted flowers occurring amongst the basal clades of our phylogeny (*Lecanthus*, *Achudemia*), followed by three or four-parted flowers (clades C1-C4) and three or two-parted flowers (clades C5-C7). This includes a clade comprising seven South Pacific species (sampled here as *Haroldiella*) characterised by two-parted female flowers (Florence, 1997). Our analysis of morphological traits recovered three-parted female flowers as plesiomorphy for the node comprising clades C5-C7 (Fig. S19). The trend in reduced female part number was matched by a reduction in male flower part number from an ancestral number of five to a derived condition of four, or three in the case of clade C1 (Fig. S15) a transition from imbricate to valvate sepals (Fig. S16). The transition from imbricate to valvate arrangement may suggest a transition from non-explosive to explosive anthesis whereby imbricate sepals open in a controlled fashion whilst valvate sepals are torn apart by the flexing filaments. The trend of reduced female and male flower part number parallels a decrease in achene size and ornamentation, *Lecanthus*, *Achudemia* and clades C1-C4 having achenes > 0.8 mm and ornamented as the ancestral state, whilst clades C5-C7 have achenes ≤ 0.8 mm and clade C7 has smooth surfaces as their ancestral states. It also parallels a change from imbricate to valvate sepals in the male flowers, suggesting a transition from gradual to explosive flower opening (Pedersoli et al., 2019).

Whilst we did not seek to test hypotheses about key innovations, we are able to use the results of our ASR analysis to propose hypotheses for future testing. Reduction in merism has been associated with key innovations in floral evolution (De Craene, 2016; Kümpers et al., 2016; Simões et al., 2017). Our study supports this, a reduction in merism coinciding with an evolutionary radiation, basal five-parted flowers of *Lecanthus* and *Achudemia* being relatively species-poor (3 spp, and 5 spp respectively). *Pilea*, in contrast, which has four-, three-, or two-parted flowers comprises ca 715 spp (Monro, 2004). According to De Craene (2016) a reduction in merism may be driven by an advantage in reducing flower size. Whilst we have not directly measured flower size, achene size is an effective surrogate for the size of the female flower and fruit, both of which are dominated by the single ovary, and later achene. As discussed above, there has been a decrease in achene size with the diversification of *Pilea*, and specifically the species-rich C7 clade. The decrease in achene size coincides with a loss of achene surface ornamentation. Based on an assumption that achene ornamentation indicates animal dispersal, our results suggest a shift in both pollen and fruit dispersal, pollen dispersal becoming explosive and more kinetic and fruit dispersal less reliant on animals.

Whilst the above discussion may provide the basis for future research into a reduction in merism and achene surface ornamentation as key innovations, the increase in species number with which they are associated may be unrelated. Increasing species number, focussed in the Greater Antilles and Andes could also be the result of increased reproductive isolation and subsequent speciation through random drift in steeply dissected shaded habitats devoid of strong air-currents. It could, therefore be, an example of an intrinsic driver of speciation rather than a response to the colonisation of novel habitats or development of novel pollination syndromes (Tilston Smith et al., 2014).

### 4.4. Proposal for an infrageneric classification

To date, there have been only two main infrageneric classifications both of which were based on morphological traits (Weddell, 1856, 1869; Chen, 1982; Chen and Monro, 2003). Whilst these classifications facilitate the identification of taxa, they are not good indicators of evolutionary relationships (Monro, 2006).

Chen (1982) proposed a classification for the Chinese taxa that focuses on the traits of female flower merism, leaf nervation, male inflorescence arrangement and male sepal arrangement. This was predated by Weddell’s classifications (Weddell, 1856, 1869) based on geographical distribution, leaf incision and heteromorphy. With the exceptions of male sepal arrangement and male inflorescence becoming receptacle-like, the traits used in both classifications were recovered as homoplastic, confirming the results of previous studies (Monro, 2006; Wu et al., 2013, 2015).

We did, however, recover achene morphology as a plesiomorphy useful in the distinction of taxa in the species-rich clade C7 from clades C1-C6 (Fig. S21). Where traits were recovered as homoplastic, they could still be used in combination, and or, with the addition of geographical distribution to delimit monophyletic infrageneric groupings. For example, clade C6, which we propose as section (§) *Plataniflora*, can be delimited by the combination of entire leaf margins, four-parted valvate male flowers and ornamented achene surface. In this way we were able to delimit seven infrageneric groupings.

The sections (§) that we propose for *Pilea* are, § *Trimeris*, § *Lecanthoides*, § *Angulata*, § *Tetrameris*, § *Verrucosa*, § *Plataniflora* and § *Leiocarpa* (Fig. 3 & Fig. 5).

## 5. Conclusions

We demonstrate that the species-rich genus *Pilea* is paraphyletic with respect to *Achudemia* and polyphyletic with respect to *Haroldiella*. We identify sepal number, flower isomery, male sepal arrangement and achene surface ornamentation as phylogenetically informative traits useful in both the delimitation of generic and infrageneric groupings and the generation of hypotheses about the evolution of the genus and the richness of its neotropical clade. Translating these findings into taxonomic actions resulted in the resurrection of *Achudemia* and the revised delimitation and infrageneric classification of *Pilea*. Our findings provide a stable framework for future research aimed at answering broader questions in evolutionary biology, such as whether intrinsic factors can drive species radiations.

## 6. Taxonomy

### 6.1. Revised delimitation of Achudemia and Pilea

**Achudemia** Blume, Mus. Bot. 2: 57, 1856. Genus type: *A. javanica* Blume, Mus. Bot. 2: 57, 1856.

*Smithiella Dunn*. Bull. Misc. Inform. 1920: 210, 1920. Genus type: *Smithiella myriantha* Dunn. nom. illeg., non *Smithiella* H. Perag. & Perag. = *Aboriella* Bennet, Indian Forester 107: 437, 1981. = *Dunniella* Rauschert, Taxon 31: 562, 1982.

Herbs, perennial or annual. Stems without stinging hairs, not releasing watery latex when cut. Leaves opposite, frequently subequal at each node, the margins toothed; cystoliths fusiform; stipules borne in axils of the leaves, persistent or caducous. Inflorescences unisexual or bisexual, paniculate, capitate or racemose cymes; pedicels subtended by inconspicuous bracteoles. Male flowers 5-merous; sepals imbricate in bud, equal, each bearing a subapical appendage. Female flowers 5-merous; the sepals 5, equal or subequal, not dimorphic. Achenes compressed ovoid, ornamented.

Five species, restricted to the Palearctic and IndoMalayan biogeographic regions. Associated with forested rocky habitats. *Achudemia japonica* is used as a medicine to treat fever and as a diuretic (Chen and Monro, 2003)

Note:— Blume (1856) in his description of the genus refers to polygamous, hermaphrodite flowers. We believe this to have been an editorial error as neither, material collected by Blume at L, or the illustration which serves as type, include polygamous or hermaphrodite flowers.

#### New combinations and typifications

***Achudemia subpubera*** (Miq.) Y.G.Wei & A.K.Monro. **comb. nov.** ≡ *Pilea subpubera* Miq. Syst. Verz. Ind. Archip. 2: 102, 1854. Type: [Indonesia], Bandong Province, *H. Zollinger 870* (holotype, U (U0226171*); isotypes P (P 02428341*), (P 02428342*))

*Achudemia javanica* Blume, Mus. Bot. 2: 57, 1856. TYPE: Mus. Bot. 2: 57, 1856, t. 20 (holotype). Epitype (selected here): [Indonesia] Java, *C.L. Blume* s.n. L (l0039782)*

We have selected an epitype as the type material comprises an illustration (https://www.biodiversitylibrary.org/item/200679#page/274/mode/1up) which is not adequate for making observations of anatomy of the leaf or stem. We selected material collected by Blume in Java, which may have served as the subject of the type illustration.

***Achudemia boniana*** (Gagnep.) L.F.Fu & Y.G.Wei. **comb. nov.** ≡ *Pilea boniana* Gagnap., Bull. Soc. Bot. France 75: 71. 1928. Type: Indochina [Vietnam], [Hà Nam Province] Kien-khé, Dong-ham rocks, *R.P. Bon* 2522 (holotype P (P06817992)*). Epitype (selected here): [Vietnam], Tonkin, Dong-Dang, on calcareous rocks, 12 Feb. 1886, B. *Balansa 581* (P (P06817995)*).

*P. morseana* Hand.-Mazz., *Symb. Sin*. 7: 140. 1929. Type: China, Guangxi, Longzhou, *Morse 495* (holotype K (K000708579)*)

*P. pentasepala* Hand.-Mazz., Symb. Sin. 7: 128. 1929. Type: China: Yunnan, mountains of Mengzi, 1800 m, *Henry 9771* (holotype K (K000708578)*)

We have selected an epitype for *Achudemia boniana* as the holotype comprises leafless material and leaves include several traits useful for species delimitation in *Achudemia*.

***Achudemia hilliana*** (Hand.-Mazz.) L.F.Fu & Y.G.Wei. **comb. nov.** ≡ *Pilea hilliana* Hand.-Mazz., Symb. Sin. 7: 129. 1929. Type: China, Yunnan, Möngdse [Mengzi], *Henry 10295* (lectotype (selected here) K (K000708583)*).

***Achudemia myriantha*** (Dunn) L.F.Fu & Y.G.Wei. **comb. nov.** ≡ *Smithiella myriantha* Dunn, Bull. Misc. Inform. Kew 1920: 211. 1920. Type: [India] Eastern Himalaya, Outer Abor Hills, sunless side of the Dihong Gorge below Rotung, 300 m. Jan 3 1912, *Burkill* 37636 (lectotype (selected here) K (K000708616)*). *Pilea myriantha* (Dunn) C.J.Chen nom. illeg., non *P. myriantha* Killip, *Bull. Bot. Res., Harbin* 2: 44. 1982. *P. spicata* C.J. Chen & A.K. Monro, Novon 17: 26. 2007.

**Pilea** Lindl., nom. cons., Coll. Bot. ad t. 4. 1821. Genus type: *P. muscosa* Lindl. nom. illeg. superfl. = *Parietaria microphylla* L. = *Pilea microphylla* (L.) Liebm.

*Adicea* Raf., nom. nud. First Cat. Gard. Transylv. Univ.: 13. 1824.

*Adicea* Raf. ex Britton & A. Br., Ill. Fl. N. U.S. 1: 533. 1896. nom. illeg. superfl.,

*Adike* Raf., New Fl. 1: 63. 1836. Genus type: *A. pumila* Raf.

*Chamaecnide* Nees & Mart. ex Miq., in C.F.P.von Martius & auct. suc. (eds.), Fl. Bras. 4: 203. 1853. Genus type: *C. microphylla* Nees ex Miq.

*Dubrueilia* Gaudich., Voy. Uranie: 495. 1830. Genus type: *D. peploides* Gaudich.

*Haroldiella* J.Florence, Fl. Polynésie Franç. 1: 218. 1997. Genus type: *H. rapaensis J.Florence*.

*Neopilea* Leandri, Ann. Mus. Colon. Marseille, sér. 6, 7-8: 46. 1950. Genus type: *N. tsaratananensis* Leandri

*Sarcopilea* Urb., Symb. Antill. 7: 201. 1912. Genus type: *S. domingensis* Urb.

Herbs, rarely shrubs, occasionally epiphytic, perennial, rarely annual. Stems without stinging hairs, not releasing watery latex when cut. Leaves opposite, frequently unequal at each node, the margins toothed or entire; cystoliths fusiform; stipules borne in axils of the leaves, persistent, rarely caducous. Inflorescences unisexual, rarely bisexual, paniculate, capitate or rarely fused cymes; pedicels subtended by inconspicuous bracteoles. Male flowers 4- or rarely 2- or 3-merous; sepals valvate, equal, each bearing a subapical appendage. Female flowers 3-, or rarely 2- or 4-merous, unequal, dimorphic, the adaxial sepal of the larger sepal frequently bearing a dorsal thickening. Achenes weakly to strongly compressed ovoid to sub-ellipsoid, smooth or ornamented. Approx. 710 spp. Cosmopolitan, except for Australia and New Zealand. A number of species cultivated as ornamentals.

#### New combinations

***Pilea australis*** L.F.Fu & A.K.Monro, *nom. nov.* Replaced name: *Haroldiella rapaensis* J.Florence, *Fl. Polynésie Franç*. 1: 220 (1997). TYPE: French Polynesia, Austral Islands, Rapa, eastern flank of Nt. Perau, 610 m, 21 Jul. 1934, *H. St. John, FR. Fosberg & J. Maireau* 15643 (holotype B1SH).

*Note:*— *Pilea australis* was created as a replacement name because a homonym, *Pilea rapensis*, has been published by Forest Brown (Brown, 1935).

***Pilea sykesii*** (J.Florence) L.F.Fu & A.K.Monro. ***comb. nov.** ≡ Haroldiella sykesii* J.Florence, Fl. Polynésie Franç. 1: 221 (1997). TYPE: French Polynesia, Austral Islands, Raivavae, Anatonu, Falaise centrale, 140 m, 10 May 1992, *J. Florence & W.R. Sykes* 11336 (holotype P (P 00637067)*, isotype PAP)

### 6.2. Key to the sections of Pilea

1. Male and female flowers with the same merism, 3-parted or 4-parted **2**

1. Male and female flowers with different merisms, 2-, 3- or 4-parted **3**

2. Merism of 3, rarely 4 (*P. tetraphylla*), stipules ≤ 10 mm in length, tropical Africa, neotropics. **§ Trimeris**

2. Merism of 4, stipules > 10 mm in length, Indomalaya. **§ Tetrameris**

3. Male inflorescence an unbranched and fused receptacle-like capitulum, involucrate.

**§ Lecanthoides**

3. Male inflorescence branched or unbranched, where unbranched capitulum globose or subglobose, not involucrate **4**

4. Achenes ornamented **5**

4. Achenes not ornamented or rarely so, where ornamented Indomalayan and either *P. melastomatoides*, or *P. peploides*. **§ Leiocarpa**

5. Stipules > 10 mm in length. **§Angulata**

5. Stipules ≤ 10 mm in length **6**

6. Leaf margins incised. **§ Verrucosa**

6. Leaf margins entire. **§ Plataniflora**

### 6.3. Infrageneric classification of Pilea

*Pilea* **§** *Trimeris* Y.G. Wei & A.K. Monro, **sect. nov.** — Section type: *P. tripartita* A.K. Monro.

Herbs. Stipules ≤ 10 mm in length. Leaf margin incised. Male inflorescence a capitate cyme, involucrate; male and female flowers with the same merism, three-rarely four-parted; the achene > 0.8 mm in length, not ornamented. Ca three spp. Tropical Africa, Neotropics.

*Pilea* **§** *Lecanthoides* C.J.Chen, Bull. Bot. Res. 2(3): 118. 1982. Section type: *P. receptacularis* C.J.Chen.

Herbs. Stipules ≤ 10 mm in length. Leaf margin incised. Male inflorescence an unbranched and fused capitulum, involucrate; male and female flowers with different merism, male flowers four-parted, female flowers three-parted; the achene > 0.8 mm in length, ornamented. Two spp. Indomalaya.

*Pilea* **§** *Angulata* L.F.Fu & Y.G.Wei, **sect. nov.** — Section type: *P. angulata* (Blume) Blume Herbs. Stipules > 10 mm in length. Leaf margin incised. Male inflorescence a branched cyme; male and female flowers with different merism, male flowers four-parted, female flowers three-parted; achene > or ≤ 0.8 mm in length, ornamented. Ca two spp. Indomalaya.

*Pilea* **§** *Tetrameris* C.J.Chen, Bull. Bot. Res. 2(3): 44. 1982. Section type: *P. basicordata* W.T.Wang.

Herbs. Stipules > 10 or rarely ≤ 10 mm in length. Leaf margin incised. Male inflorescence a branched cyme; male and female flowers with the same merism, flowers four-parted; achene > 0.8 mm in length, ornamented. Ca 15 spp. Indomalaya.

*Pilea* **§** *Verrucosa* L.F.Fu & Y.G.Wei, sect. nov. Type: *P. gracilis* Hand.-Mazz.

Herbs. Stipules ≤ 10 mm in length. Leaf margin incised. Male inflorescence a branched cyme; male and female flowers with different merism, male flowers four- or occasionally two-parted, female flowers three-parted, rarely two-parted; achene > or ≤ 0.8 mm in length, ornamented. Ca 80 spp. Indomalaya.

*Pilea* **§** *Plataniflora* L.F.Fu & Y.G.Wei, sect. nov. Type: *P. plataniflora* C.H.Wright.

Herbs. Stipules ≤ 10 mm in length. Leaf margins entire. Male inflorescence a branched cyme; male and female flowers with different merism, male flowers four-parted, female flowers three-parted; achene > or ≤ 0.8 mm in length, ornamented. Ca 34 spp. Indomalaya.

*Pilea* **§** *Leiocarpa* L.F.Fu & Y.G.Wei, sect. nov. Type: *P. micropylla* (L.) Liebm.

Herbs. Stipules > or ≤ 10 mm in length. Leaf margins entire or incised. Male inflorescence a branched or capitate cyme; male and female flowers with different merism, male flowers four- or rarely two-parted, female flowers four- or rarely two-parted, achene > or ≤ 0.8 mm in length, not ornamented. Ca 570 spp, Indomalaya, Neotropics, Australasia, Palearctic, Nearctic.

### 6.4. Excluded names

*Metapilea* W.T.Wang, Bull. Bot. Res., Harbin 36: 164. 2016. Genus type: *M. jingxiensis* W.T.Wang.

We have some doubt over the position of this taxon based on the poor quality of the material upon which the description and illustration are based (a single sheet, very few immature flowers). One individual of this taxon was collected once in a relatively common habitat in Guangxi Province, China, in 1973 (*Wang* s.n.) and it has not been collected since. It is possible that the original material may be an immature collection of *Pilea*, to which it is vegetatively identical, or a distinct genus, which, based on the illustrations, could be allied to *Elatostema* or *Procris*. Due to the sampling policy of the herbarium where the type collection is stored it is not possible to sample this material for DNA and so its status and position remains uncertain.

## Supporting information

combined supplementary materials

## Acknowledgements

We would like to thank Robert von Blittersdorff and David Schellenberger Costa (African Plants - A photo guide) for high resolution images of two *Pilea* species, Nicholas Hind (K) for help with nomenclatural queries and Margaret Tebbs (K) for the illustrations, Guo-Xiong Hu (GZU) and Zi-Bing Xin (GXIB) for joining fieldtrips. We are grateful to K, KUN and SING for providing DNA samples of some key taxa. This work was supported by the National Natural Science Foundation of China (grant number 31860042), Guangxi Natural Science Foundation Program (grant number 2017GXNSFBA198014), and the STS Program of the Chinese Academy of Sciences (grant number KFJ-3W-No1).

## REFERENCES

Bennet, S.S.R., Raizada, M.B., 1981. Nomenclatural changes in some flowering plants. Indian Forester 107, 432–437.

Blume, C.L., 1856. Museum botanicum Lugduno-Batavum: 2. Vol. 2. Brill, Leiden, pp. 57.

Brizicky, G.K., 1969. Subgeneric and sectional names: their starting points and early sources. Taxon 18, 643–660.

Brown, F.B.H., 1935. Flora of southeastern Polynesia, III. Dicotyledons. Bishop Mus. Bull. 130, 386.

Chen, C.J., 1982. A monograph of *Pilea* (Urticaceae) in China. Bull. Bot. Res. Harbin 2, 1–132.

Chen, C.J., 1995. *Pilea*, in: Wang, W.T. and Chen, C.J. (Eds.), Flora Reipublicae Popularis Sinicae. Science Press, Beijing, pp. 57–156.

Chen, C.J., Monro, A.K., 2003. *Pilea*, in: Wu, Z.Y., Raven, P.H. (Eds.), Flora of China. Science Press and Missouri Botanical Garden Press, Beijing and St. Louis, pp.76–189.

Chen, L.Y., Song, M.S., Zhu, H.G., Li, Z.M., 2014. A modified protocol for plant genome DNA extraction. Plant Diversity Resour. 36, 375–380.

De Craene, L.R., 2016. Meristic changes in flowering plants: How flowers play with numbers. Flora 221, 22–37. https://doi.org/10.1016/j.flora.2015.08.005.

Dunn, S.T., 1920. Bulletin of Miscellaneous Information, Royal Gardens Kew, London.

Fay, M.F., Swensen, S.M., Chase, M.W., 1997. Taxonomic affinities of *Medusagyne oppositifolia* (Medusagynaceae). Kew Bull. 52, 111–120. https://doi.org/10.2307/4117844.

sFlorence, J., 1997. Flore de la Polynésie Française. Institut de recherche pour le développement, Paris.

Friis, I., 1989. Urticacea, in: Polhill, R.M. (Ed.), Flora of Topical East Africa. A.A. Balkema, Rotterdam, pp. 1–64.

Fu, L.F., Huang, S.L., Monro, A.K., Liu, Y., Wen, F., Wei, Y.G., 2017a. *Pilea nonggangensis* (Urticaceae), a new species from Guangxi, China. Phytotaxa 313, 130–136. https://doi.org/10.11646/phytotaxa.313.1.9.

Fu, L.F., Su, L.Y., Mallik, A., Wen, F., Wei, Y.G., 2017b. Cytology and sexuality of 11 species of *Elatostema* (Urticaceae) in limestone karsts suggests that apomixis is a recurring phenomenon. Nord. J. Bot. 35, 251–256. https://doi.org/10.1111/njb.01281.

Gaudichaud, C., 1830. Botanique, part 12, in: Freycinet, H.d. (Ed.), Voyage autour du monde…executé sur les corvettes de S.M. l’ Uranie et la Physiciene’. Pilet-Aine, Paris, pp. 465–522.

Habib, S., Dang, V.C., Ickert-Bond, S.M., Zhang, J.L., Lu, L.M., Wen, J., Chen, Z.D., 2017. Robust phylogeny of *Tetrastigma* (Vitaceae) based on ten plastid DNA regions: implications for infrageneric classification and seed character evolution. Front. Plant Sci. 8, 590. https://doi.org/10.3389/fpls.2017.00590.

Hao, Z., Kuang, Y., Kang, M., 2015. Untangling the influence of phylogeny, soil and climate on leaf element concentrations in a biodiversity hotspot. Funct. Ecol. 29, 165–176. https://doi.org/10.1111/1365-2435.12344.

Huelsenbeck, J.P., Ronquist, F., 2001. MRBAYES: Bayesian inference of phylogenetic trees. Bioinformatics 17, 754–755.

IPNI, 2020. International Plant Names Index. http://www.ipni.org/ (accessed 26 May 2020)

Jestrow, B., Valdés, J.J., Jimenez Rodriguez, F., Francisco-Ortega, J., 2012. Phylogenetic placement of the Dominican Republic endemic genus *Sarcopilea* (Urticaceae). Taxon 61, 592–600. https://doi.org/10.1002/tax.613008.

Katoh, K., Standley, D.M., 2013. MAFFT multiple sequence alignment software version 7: improvements in performance and usability. Mol. Biol. Evol. 30, 772–780. https://doi.org/10.1093/molbev/mst010.

Kim, C., Deng, T., Chase, M., Zhang, D. G., Nie, Z.L., Sun, H., 2015. Generic phylogeny and character evolution in Urticeae (Urticaceae) inferred from nuclear and plastid DNA regions. Taxon 64, 65–78. https://doi.org/10.12705/641.20.

Kümpers, B.M.C., Richardson, J.E., Anderberg, A.A., Wilkie, P., Ronse De Craene, L.P., 2016. The significance of meristic changes in the flowers of Sapotaceae. Bot. J. Linn. Soc. 180, 161–192. https://doi.org/10.1111/boj.12363.

Maddison, W.P., Maddison, D.R., 2015. Mesquite: a modular system for evolutionary analysis. Version: 3.51 ed.

Miller, M.A., Pfeiffer, W., Schwartz, T., 2010. Creating the CIPRES Science Gateway for inference of large phylogenetic trees. Proceedings of the Gateway Computing Environments Workshop (GCE), 2010. Institute of Electrical and Electronics Engineers, New Orleans, 1–8.

Monro, A.K., 1999. Seven new species of *Pilea* Lindley (Urticaceae) from Mesoamerica. Novon 9, 390–400. https://doi.org/10.2307/3391738.

Monro, A.K., 2004. Three new species, and three new names in *Pilea* (Urticaceae) from New Guinea. Kew Bull. 59, 573–579. https://doi.org/10.2307/4110914.

Monro, A.K., 2006. The revision of species-rich genera: a phylogenetic framework for the strategic revision of *Pilea* (Urticaceae) based on cpDNA, nrDNA and morphology. Am. J. Bot. 93, 426–441. https://doi.org/10.3732/ajb.93.3.426.

Monro, A.K., 2015. Urticaceae, in Davidse, G., Sousa Sanchez, M., Knapp, S., Chiang Cabrera, F. (Eds) Saururaceae a Zygophyllaceae, Flora Mesoamericana. Missouri Botanical Garden, Saint Louis, pp. 116–174.

Monro, A.K., Wei, Y.G., Chen, C.J., 2012. Three new species of *Pilea* (Urticaceae) from limestone karst in China. Phytokeys 19: 51–66. https://doi.org/10.3897/phytokeys.19.3968.

Pedersoli, G.D., Leme, F.M., Leite, V.G., Teixeira, S. P., 2019. Anatomy solves the puzzle of explosive pollen release in wind-pollinated urticalean rosids. Am. J. Bot. 106, 489–506. https://doi.org/10.1002/ajb2.1254.

Planta, V., 2003. Phylogenetic relationships of the Afro-Malagasy members of the large genus *Begonia* inferred from *trnL* intron sequences. Syst. Bot. 28, 693–704. https://doi.org/10.1043/02-56.1.

Posada, D., Crandall, K.A., 1998. ModelTest: Testing the model of DNA substitution. Bioinformatics (Oxford, England) 14, 817–818. https://doi.org/10.1093/bioinformatics/14.9.817.

Simões, M., Breitkreuz, L., Alvarado, M., Baca, S., Cooper, J.C., Heins, L., Herzog, K., Lieberman, B.S., 2017. The Evolving Theory of Evolutionary Radiations. Trends Ecol. Evol. 31, 27–34. https://doi.org/10.1016/j.tree.2015.10.007.

Stamatakis, A., Hoover, P., Rougemont, J., 2008. A rapid bootstrap algorithm for the RAxML web servers. Syst. Biol. 57, 758–771. https://doi.org/10.1080/10635150802429642.

Swofford, D.L., 2002. PAUP*: Phylogenetic analysis using parsimony (*and other methods). Sinauer Associates Inc., Sunderland, MA, USA.

Taberlet, P., Gielly, L., Pautou, G., Bouvet J., 1991. Universal primers for amplification of three non-coding regions of chloroplast DNA. Plant Mol. Biol. 17, 1105–1109. https://doi.org/10.1007/BF00037152.

Tamura, K., Peterson, D., Peterson, N., Stecher, G., Nei, M., Kumar, S., 2011. MEGA5: molecular evolutionary genetics analysis using maximum likelihood, evolutionary distance, and maximum parsimony methods. Mol. Biol. Evol. 28, 2731–2739. https://doi.org/10.1093/molbev/msr121.

Tilston Smith, B., McCormack, J.E., Cuervo, A.M., Hickerson, M.J., Aleixo, A., Cadena, C. D., Pérez-Emán J., Burney C.W., Xie X, Harvey M.G., Faircloth, B.C., Glenn T.C., Derryberry E.P., Prejean J., Fields S., Brumfield R.T., 2014. The drivers of tropical speciation. Nature 515, 406–409. https://doi.org/10.1038/nature13687.

Tseng, Y.H., Monro, A.K., Wei, Y.G., Hu, J.M., 2019. Molecular phylogeny and morphology of *Elatostema* s.l. (Urticaceae): implications for inter- and infrageneric classification. Mol. Phylogenet. Evol. 132, 251–264. https://doi.org/10.1016/j.ympev.2018.11.016.

Turland, N.J., Wiersema, J.H., Barrie, F.R., Greuter, W., Hawksworth, D.L., Herendeen, P.S., Knapp, S., Kusber, W.H., Li, D.Z., Marhold, K., May, T.W., McNeill, J., Monro, A.M., Prado, J., Price, M.J., Smith, G.F., 2018. International Code of Nomenclature for algae, fungi, and plants (Shenzhen Code) adopted by the Nineteenth International Botanical Congress Shenzhen, China, July 2017. Regnum Vegetabile 159. Glashütten: Koeltz Botanical Books.

WCVP, 2020. World Checklist of Vascular Plants, version 2.0. http://wcvp.science.kew.org/ (accessed 26 May 2020).

Weddell, H.A., 1856. Monographie de la famille des Urticées. Archives Mus. Hist. Nat. Paris. 9, 1–400.

Weddell, H.A., 1869. Urticaceae, in: De Candolle, A. (Ed.), Prodomus Systematis naturalis regni vegetabilis. Masson, Paris, pp. 32–235.

Wen, J., Nie, Z.L., Ickert-Bond, S.M., 2016. Intercontinental disjunctions between eastern Asia and western North America in vascular plants highlight the biogeographic importance of the Bering land bridge from late Cretaceous to Neogene. J. Syst. Evol. 54, 469–490. https://doi.org/10.1111/jse.12222.

White, T.J., Bruns, T., Lee, S.J.W.T., Taylor, J., 1990. Amplification and direct sequencing of fungal ribosomal RNA genes for phylogenetics. PCR protocols: a guide to methods and applications 18, 315–322.

Wu, Z.Y., Liu, J., Provan, J., Wang, H., Chen, C.J., Cadotte, M.W., Luo, Y.H., Amorim, B.S., Li, D.Z., Milne, R.I., 2018. Testing Darwin’s transoceanic dispersal hypothesis for the inland nettle family (Urticaceae). Ecol. Lett. 21, 1515–1529. https://doi.org/10.1111/ele.13132.

Wu, Z.Y., Milne, R.I., Chen, C.J., Liu, J., Wang, H., Li, D.Z., 2015. Ancestral state reconstruction reveals rampant homoplasy of diagnostic morphological characters in Urticaceae, conflicting with current classification schemes. Plos One 10, e0141821. https://doi.org/10.1371/journal.pone.0141821.

Wu, Z.Y., Monro, A.K., Milne, R.I., Wang, H., Yi, T.S., Liu, J., Li, D.Z., 2013. Molecular phylogeny of the nettle family (Urticaceae) inferred from multiple loci of three genomes and extensive generic sampling. Mol. Phylogenet. Evol. 69, 814–827. https://doi.org/10.1016/j.ympev.2013.06.022.

Yang, F., Wang, Y.H., Qiao, D., Wang, H.C., 2018. *Pilea weimingii* (Urticaceae), a new species from Yunnan, southwest China. Ann. Bot. Fenn. 55, 99–103. https://doi.org/10.5735/085.055.0112.

Zhang, S.D., Soltis, D.E., Yang, Y., Li, D.Z., Yi, T.S., 2011. Multi-gene analysis provides a well-supported phylogeny of Rosales. Mol. Phylogenet. Evol. 60, 21–28. https://doi.org/10.1016/j.ympev.2011.04.008.

