## Supplementary material for "Phylogeny of the species-rich *Pilea* Lindl. (Urticaceae) supports its revised delimitation and infrageneric classification, including the resurrection of *Achudemia* Blume": combined supplementary materials

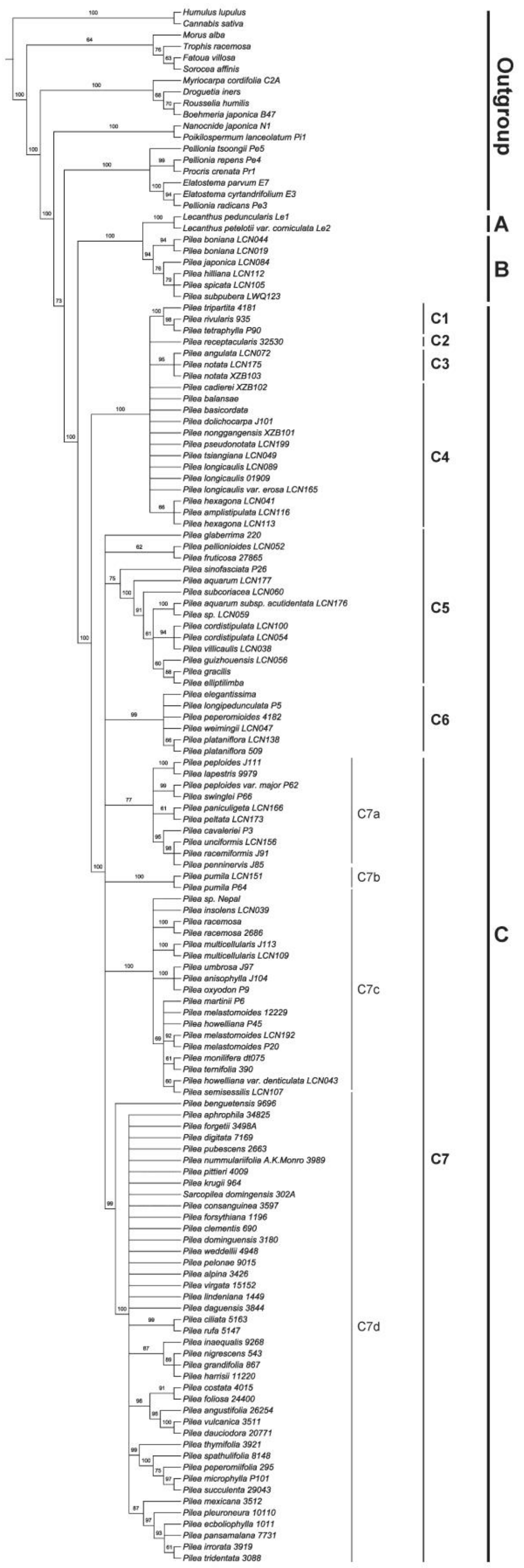


Fig. S1. Phylogenetic tree of *Pilea* generated from maximum likelihood (ML) of cpDNA dataset (*trnL-F* spacer and *rbcL*). Numbers on the branches indicate the bootstrap values (≥60%).

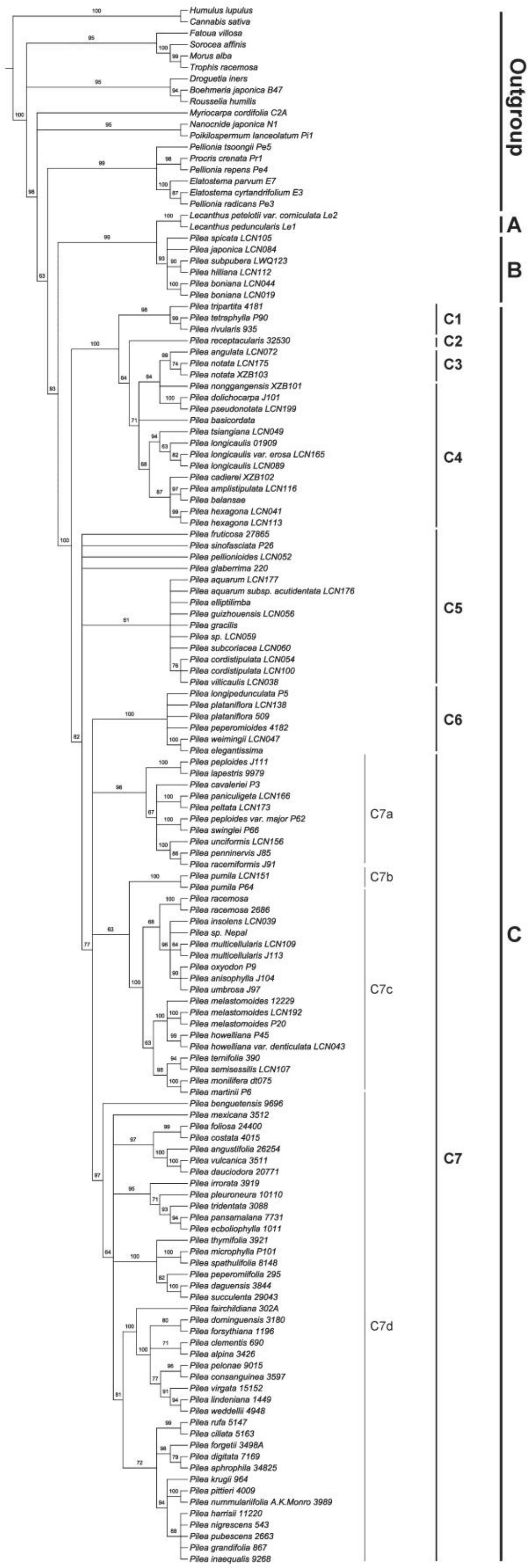


Fig. S2. Phylogenetic tree of *Pilea* generated from maximum likelihood (ML) of nrDNA dataset (nrITS). Numbers on the branches indicate the bootstrap values (≥60%).

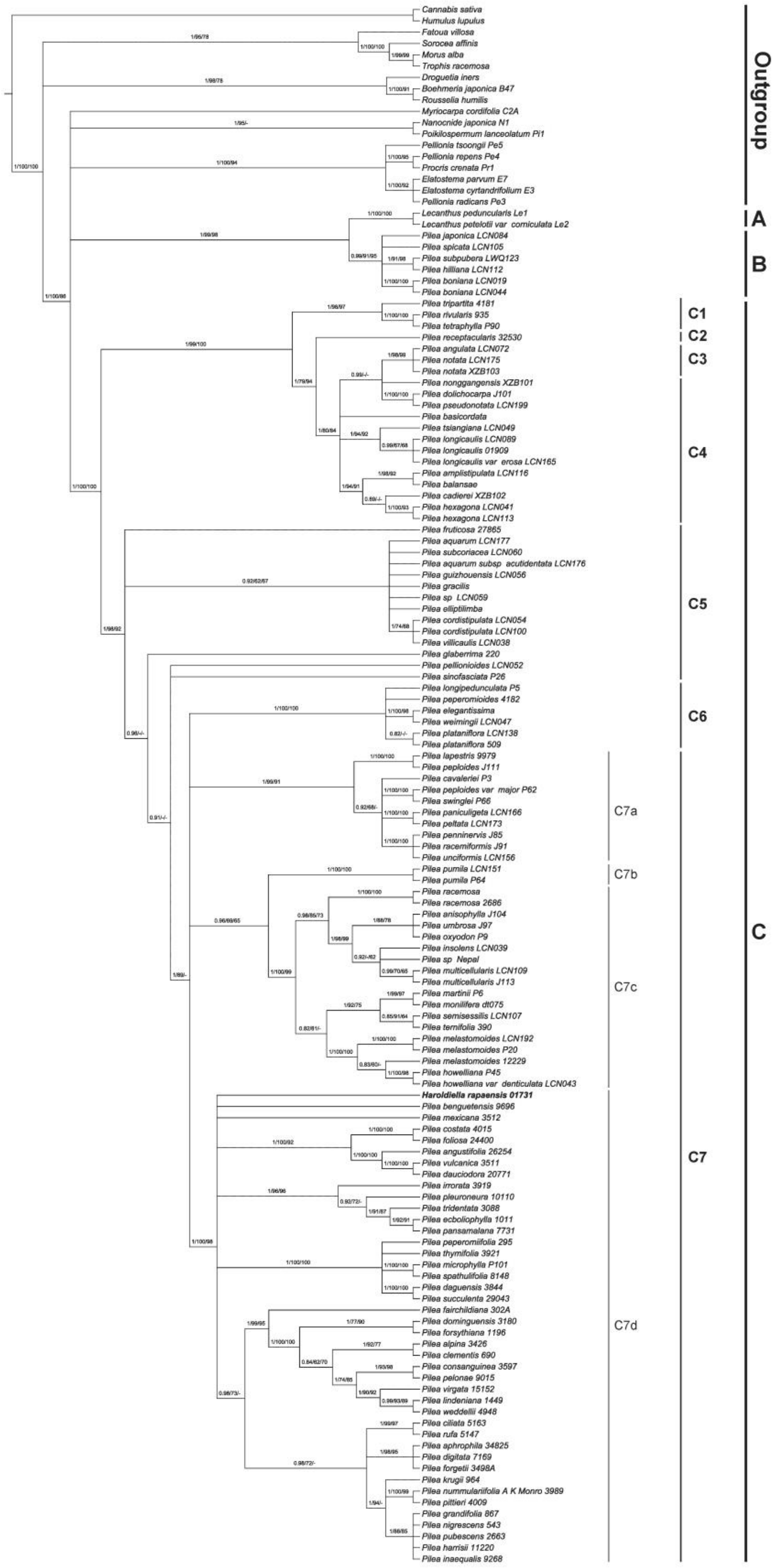


Fig. S3. Phylogenetic tree of *Pilea* (including *Haroldiella*) generated from Bayesian Inference (BI) of nrITS dataset. Numbers on the branches indicate the posterior probability (≥0.8) of BI and bootstrap values (≥60%) of the maximum likelihood (ML) and the maximum parsimony (MP) analyses.

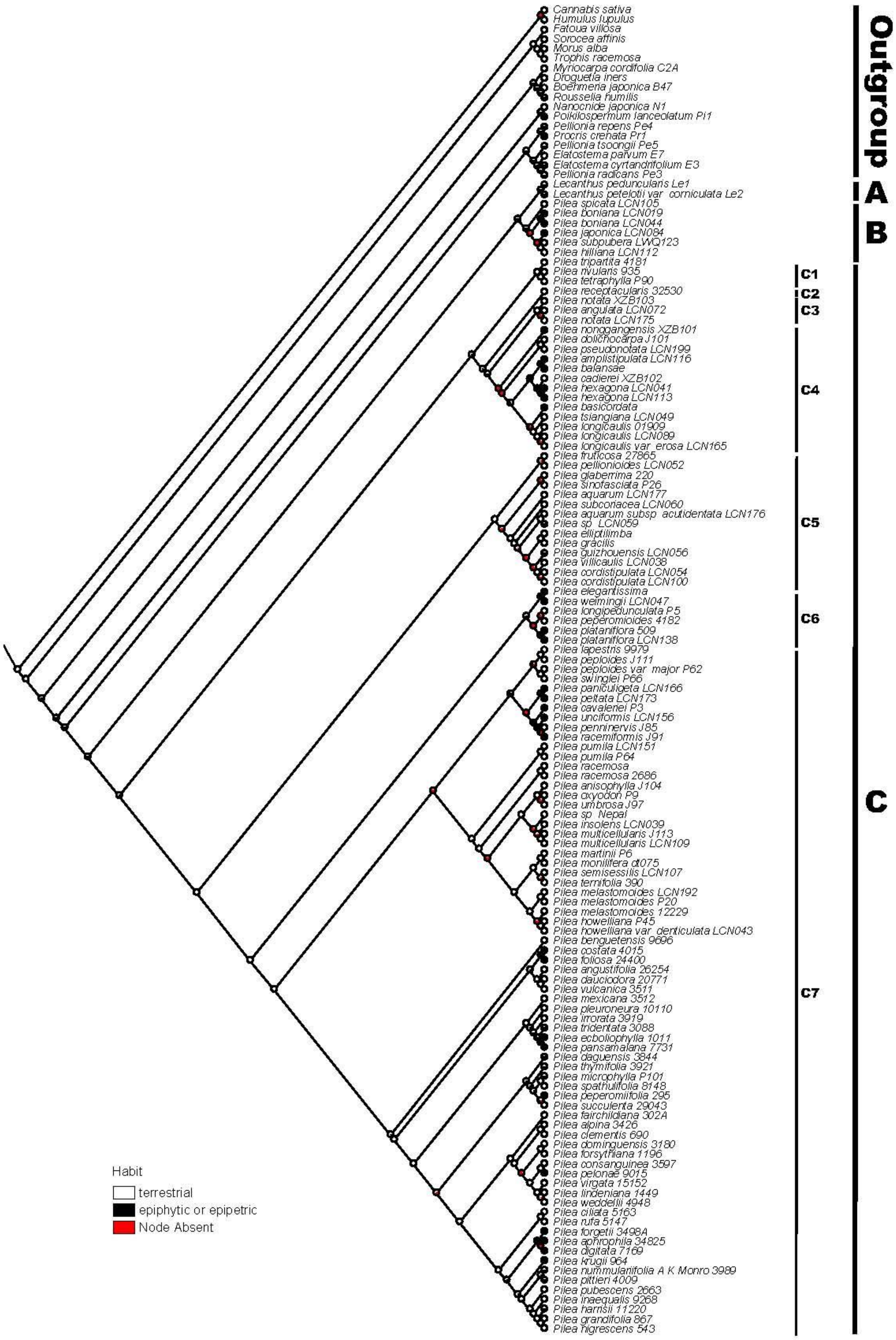


Fig. S4. Ancestral state reconstruction for *Pilea* based on Maximum likelihood analysis of habit.

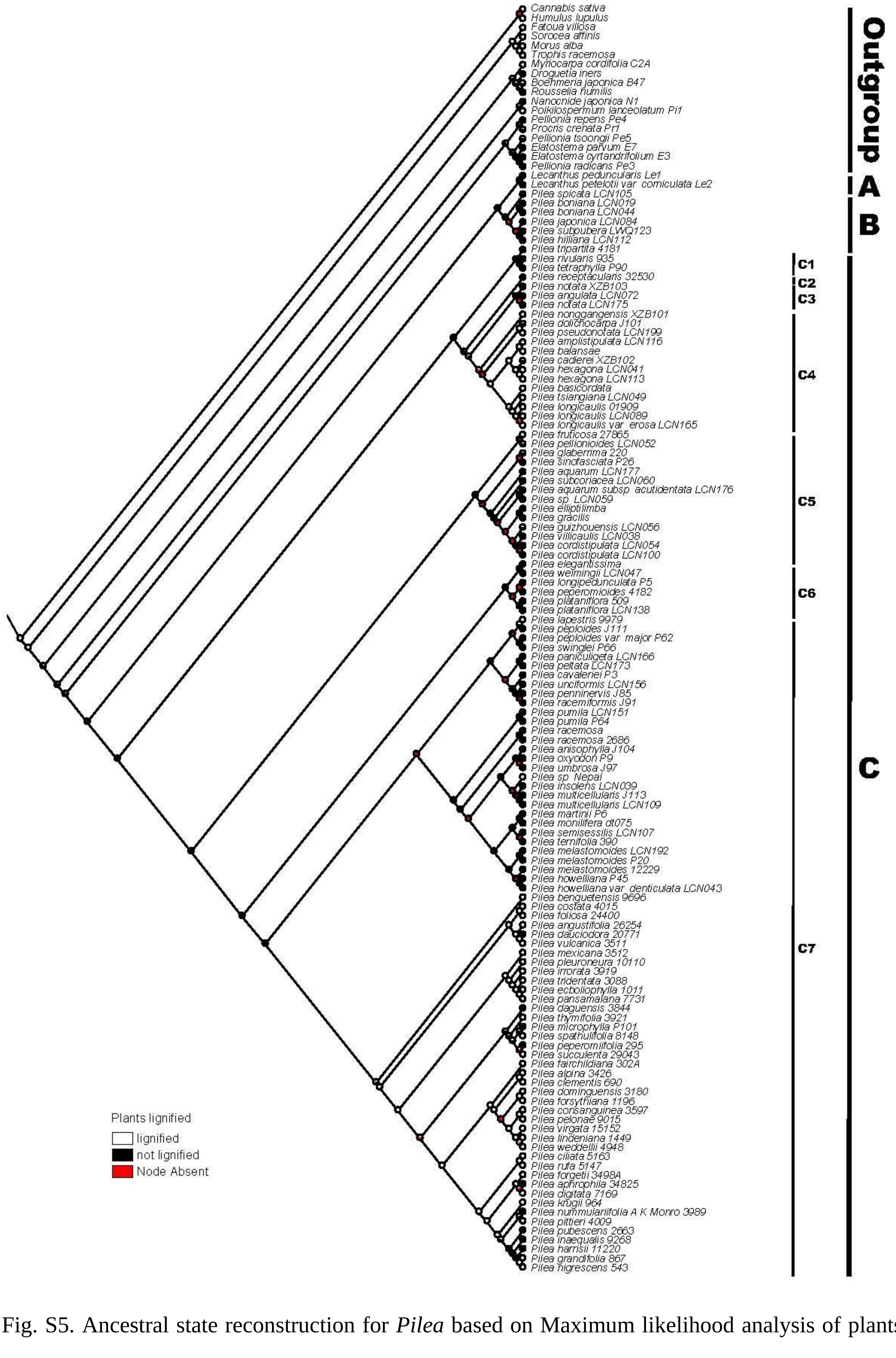


Fig. S5. Ancestral state reconstruction for *Pilea* based on Maximum likelihood analysis of plants lignified.

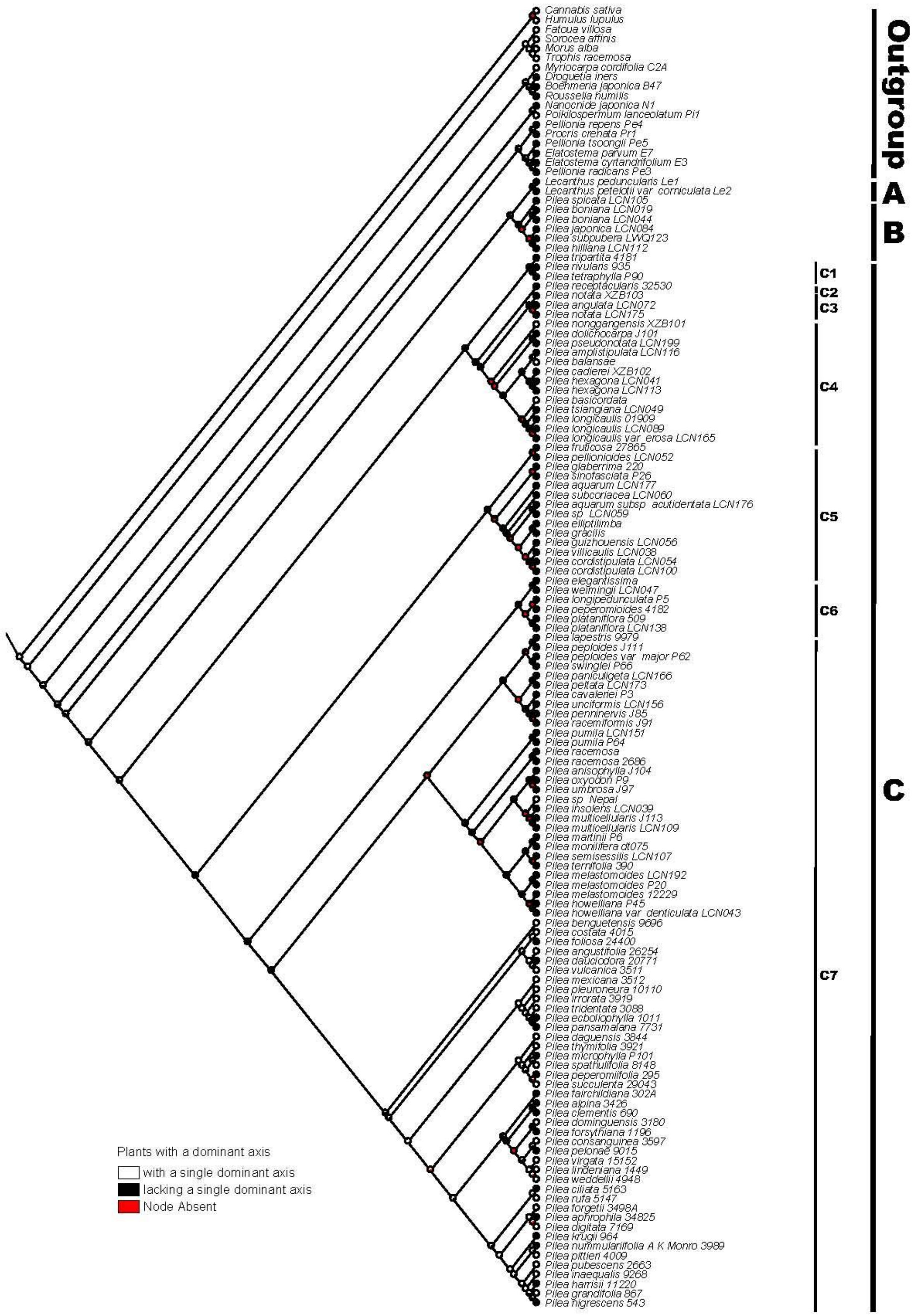


Fig. S6. Ancestral state reconstruction for *Pilea* based on Maximum likelihood analysis of plants with a dominant axis.

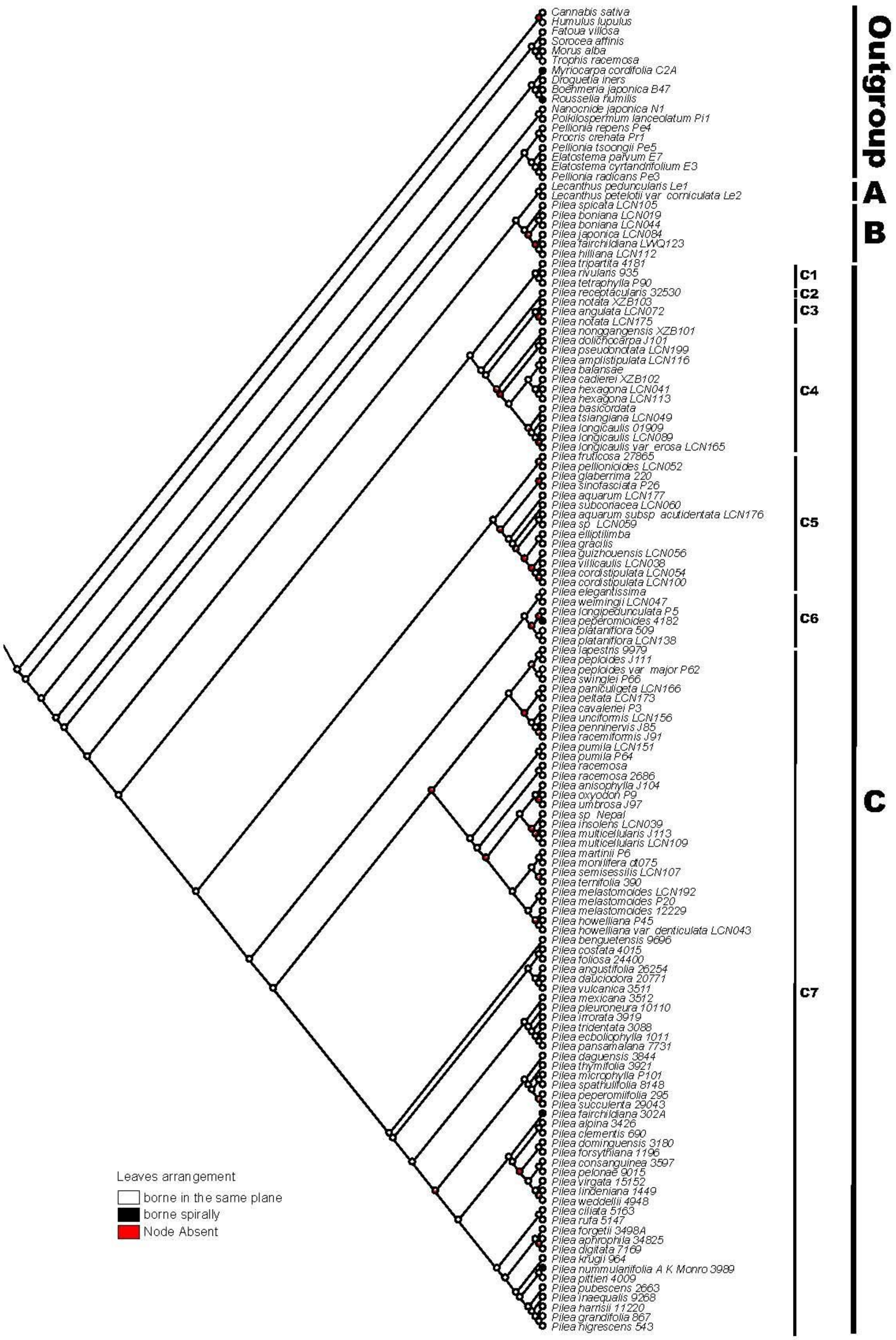


Fig. S7. Ancestral state reconstruction for *Pilea* based on Maximum likelihood analysis of leaves arrangement.

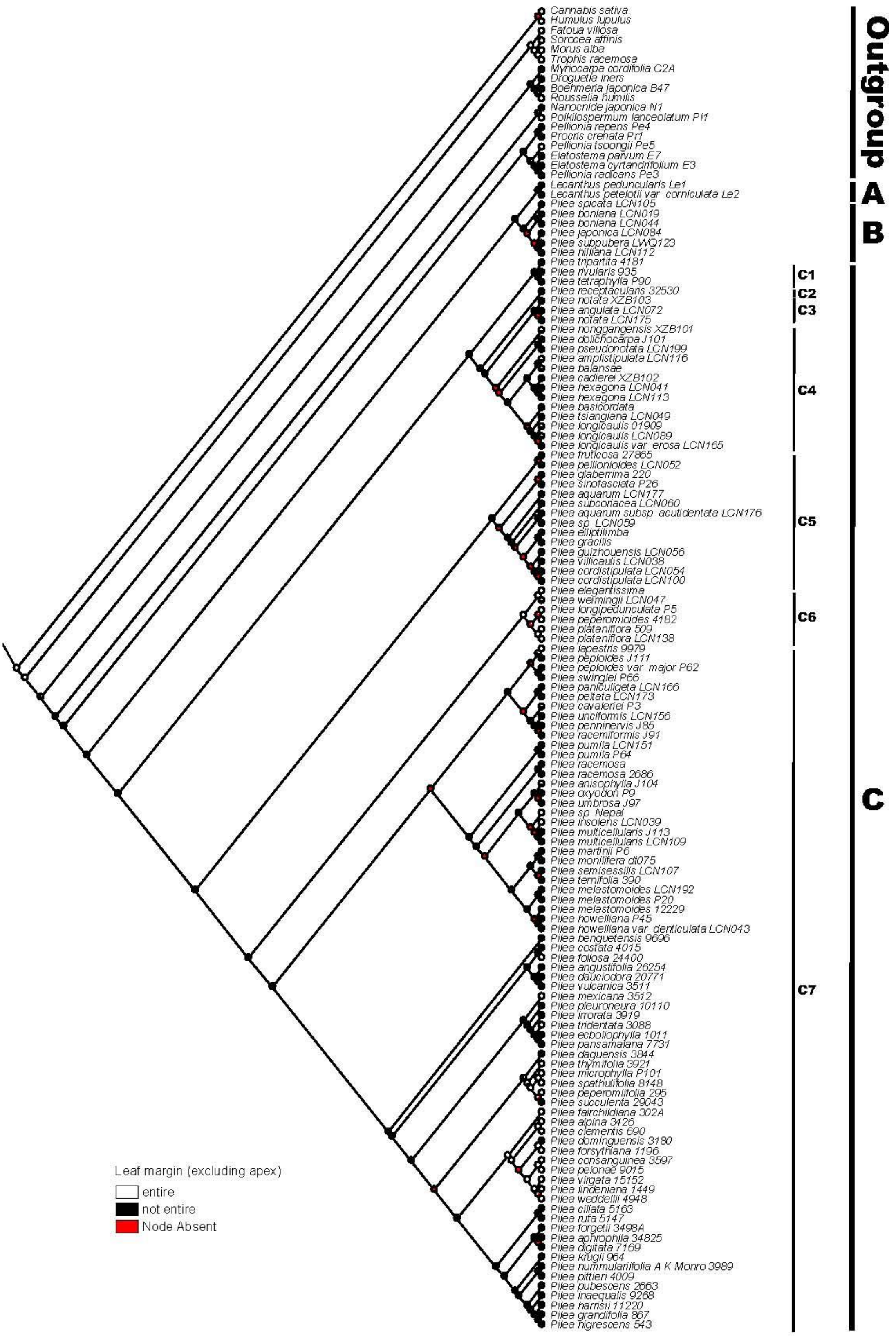


Fig. S8. Ancestral state reconstruction for *Pilea* based on Maximum likelihood analysis of leaf margin (excluding apex).

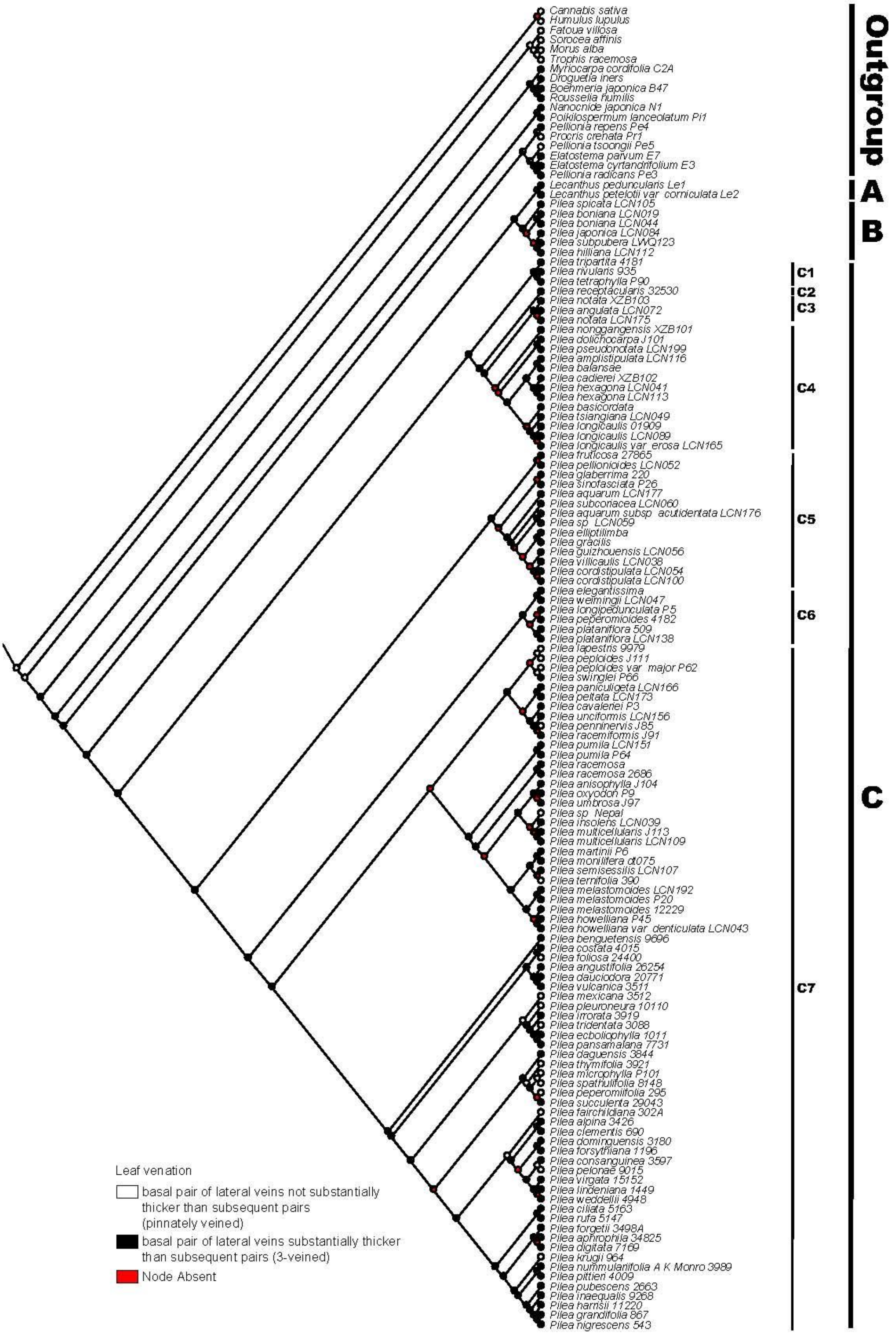


Fig. S9. Ancestral state reconstruction for *Pilea* based on Maximum likelihood analysis of leaf venation.

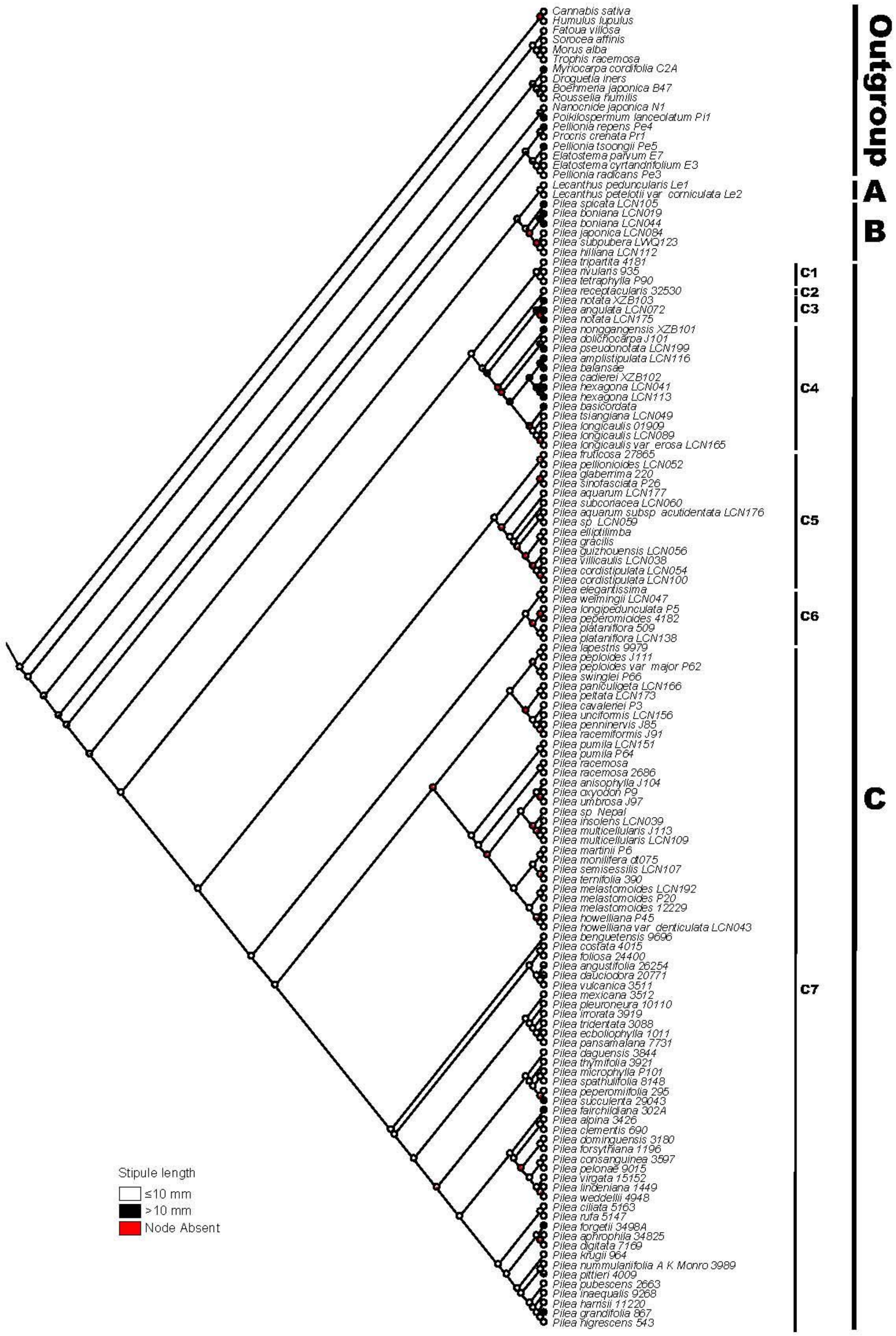


Fig. S10. Ancestral state reconstruction for *Pilea* based on Maximum likelihood analysis of stipule length.

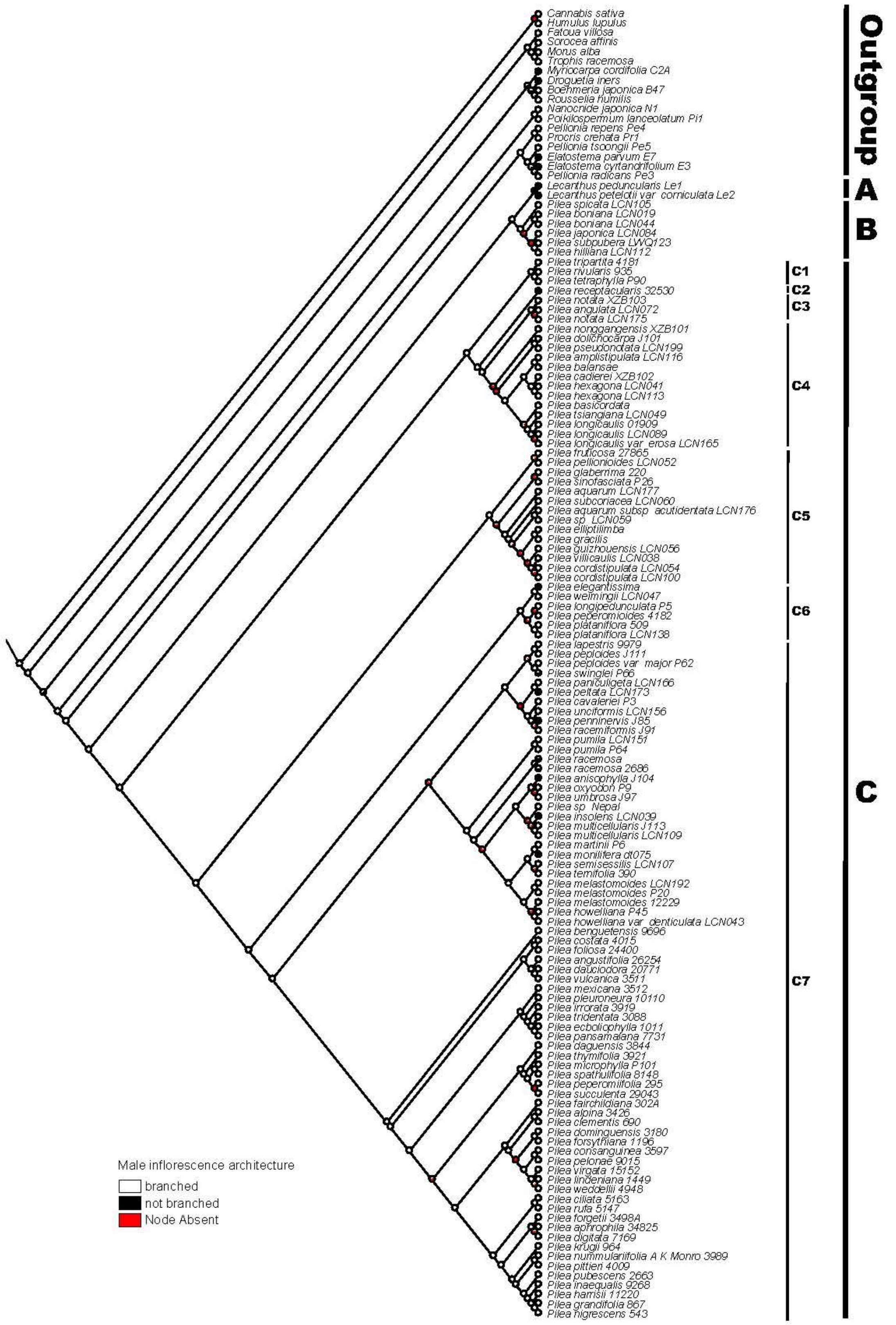


Fig. S11. Ancestral state reconstruction for *Pilea* based on Maximum likelihood analysis of male inflorescence architecture.

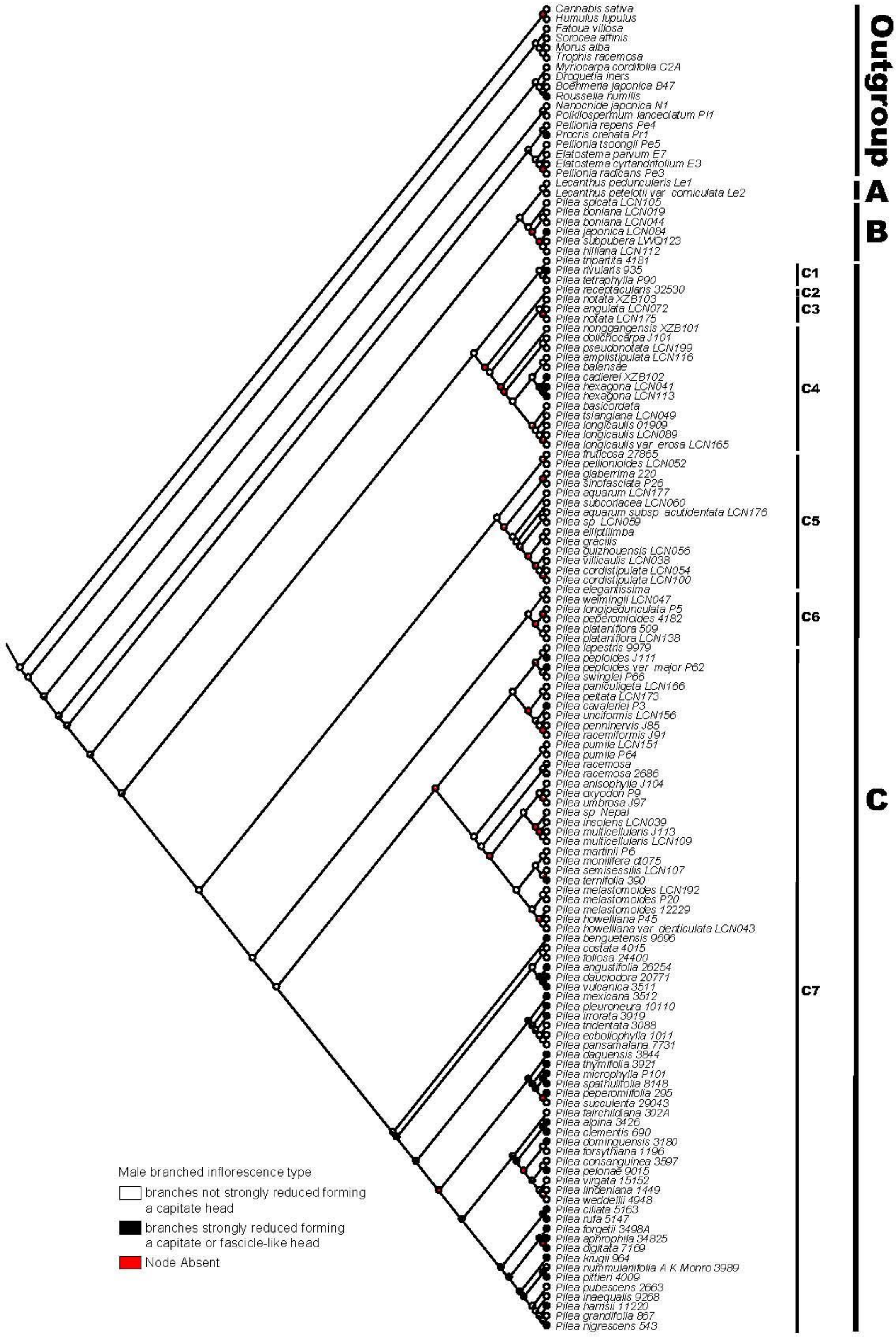


Fig. S12. Ancestral state reconstruction for *Pilea* based on Maximum likelihood analysis of male branched inflorescence type.

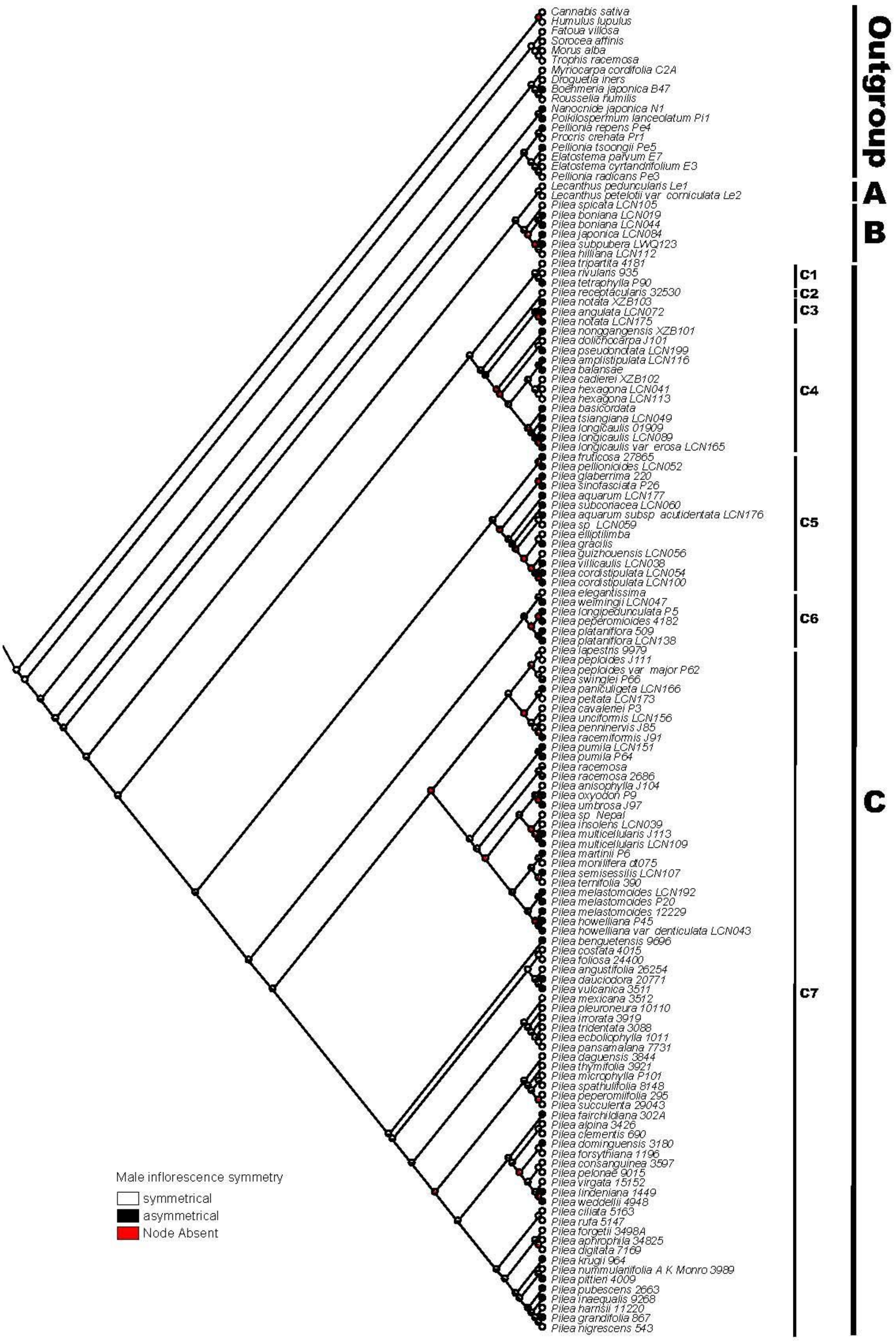


Fig. S13. Ancestral state reconstruction for *Pilea* based on Maximum likelihood analysis of male inflorescence symmetry.

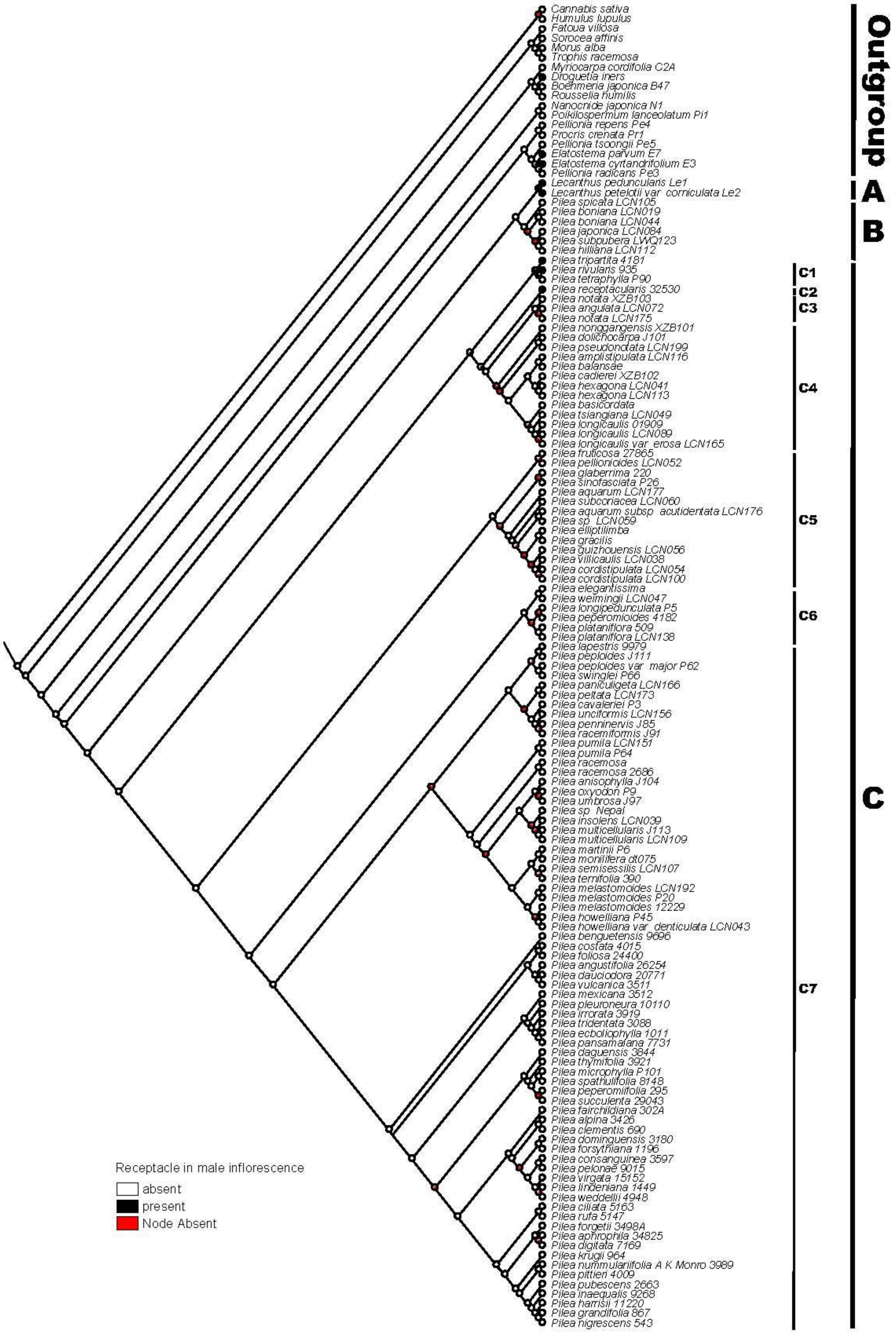


Fig. S14. Ancestral state reconstruction for *Pilea* based on Maximum likelihood analysis of receptacle in male inflorescence.

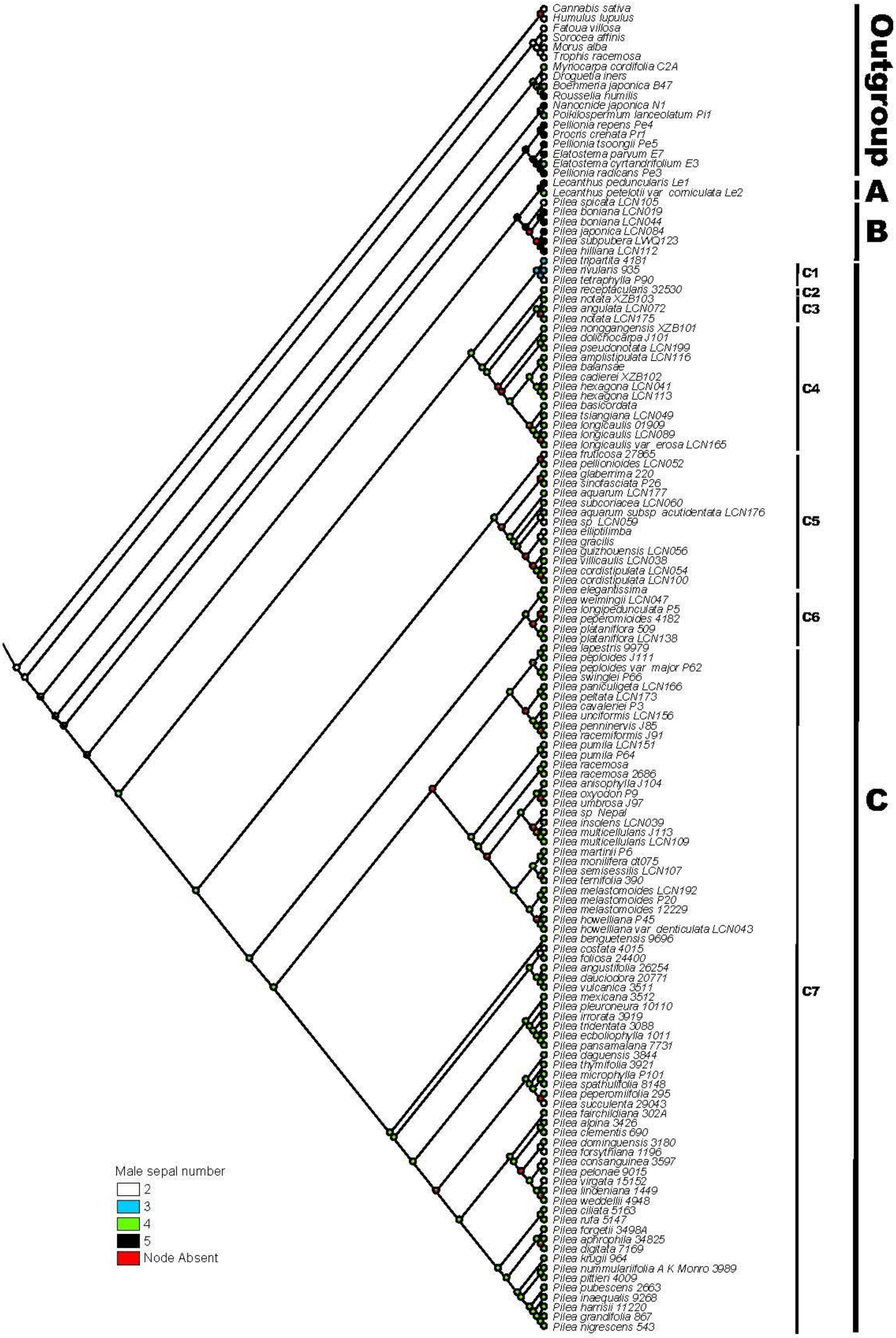


Fig. S15. Ancestral state reconstruction for *Pilea* based on Maximum likelihood analysis of male sepal number.

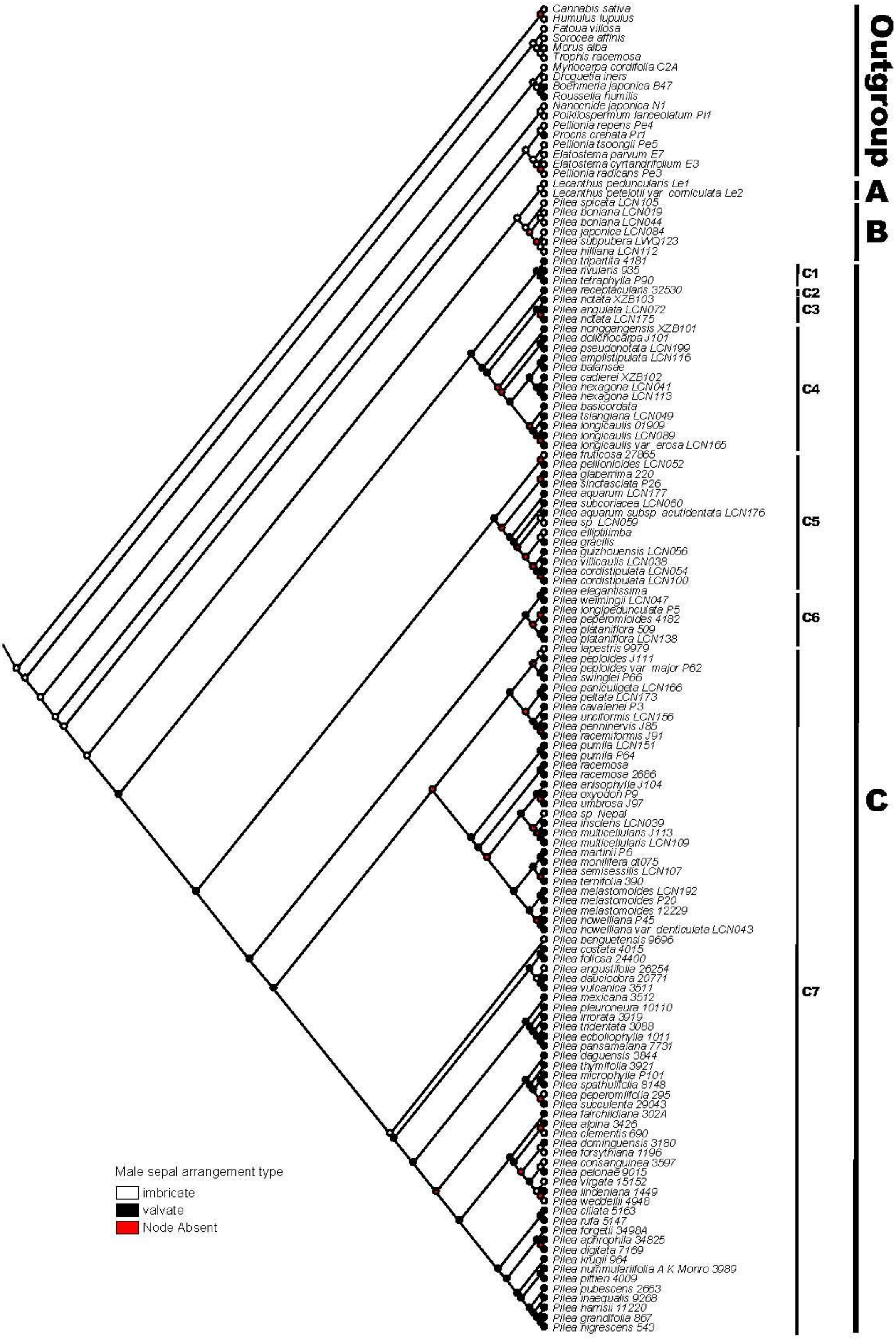


Fig. S16. Ancestral state reconstruction for *Pilea* based on Maximum likelihood analysis of male sepal arrangement type.

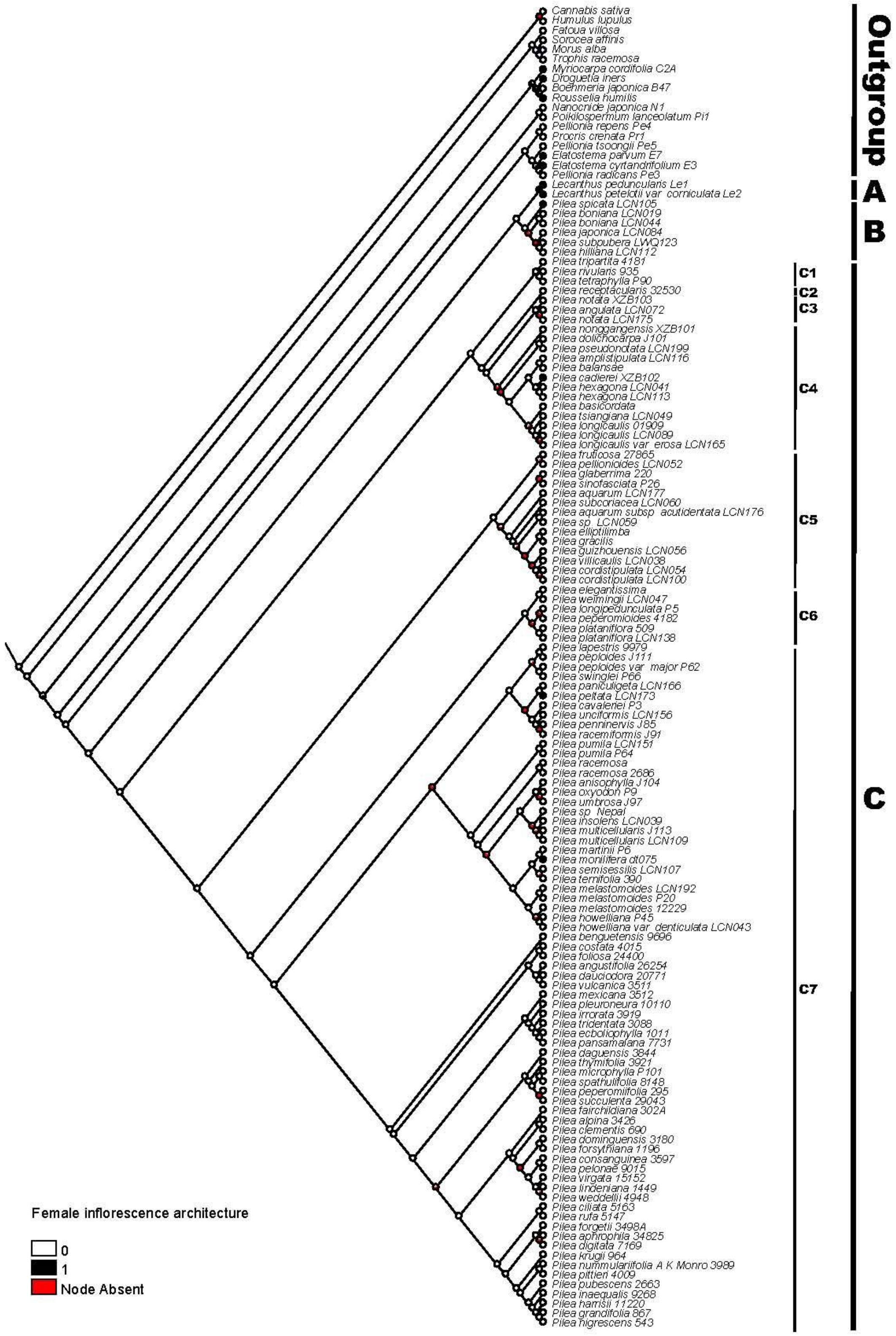


Fig. S17. Ancestral state reconstruction for *Pilea* based on Maximum likelihood analysis of female inflorescence architecture.

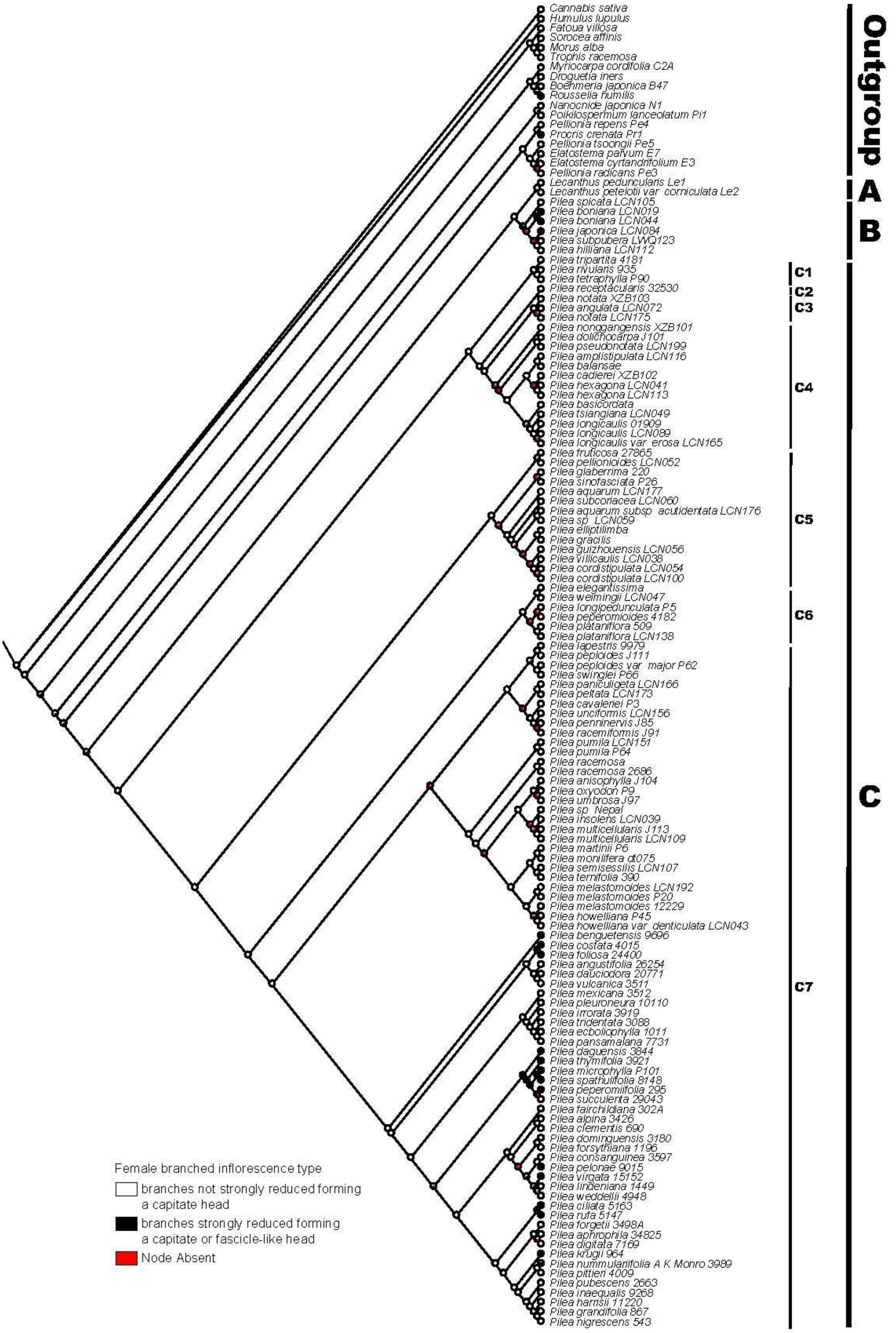


Fig. S18. Ancestral state reconstruction for *Pilea* based on Maximum likelihood analysis of female branched inflorescence type.

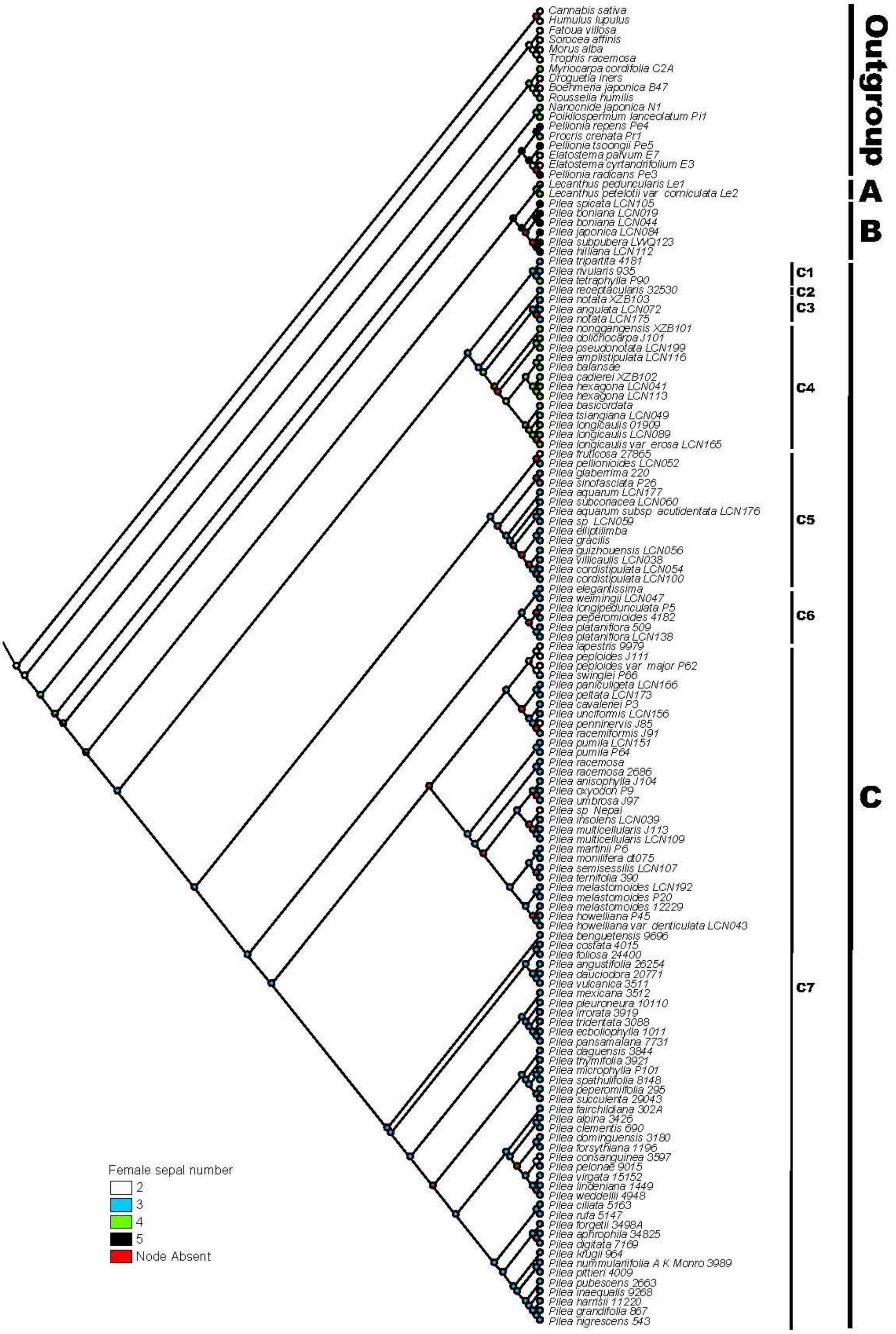


Fig. S19. Ancestral state reconstruction for *Pilea* based on Maximum likelihood analysis of female sepal number.

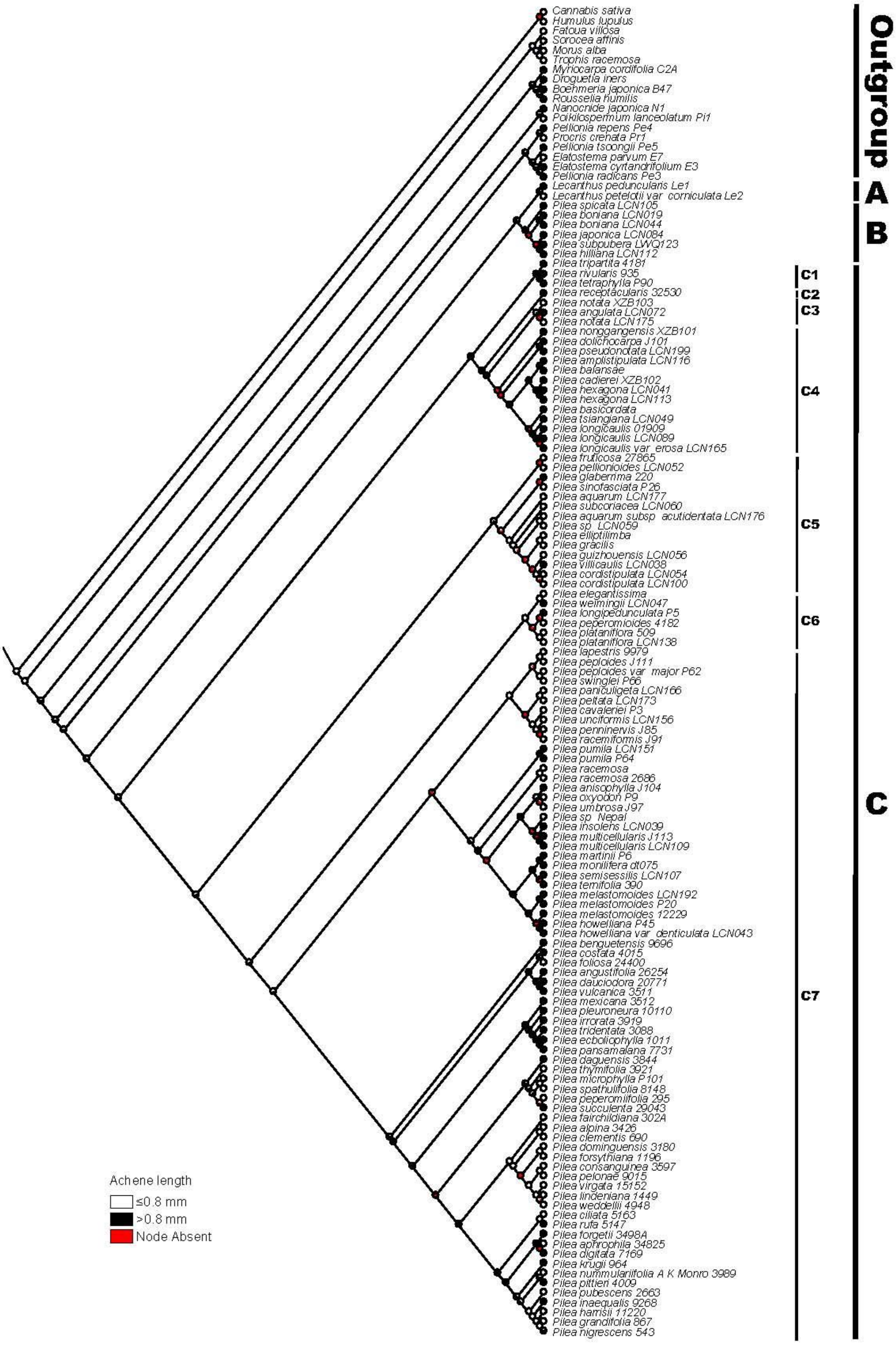


Fig. S20. Ancestral state reconstruction for *Pilea* based on Maximum likelihood analysis of achene length.

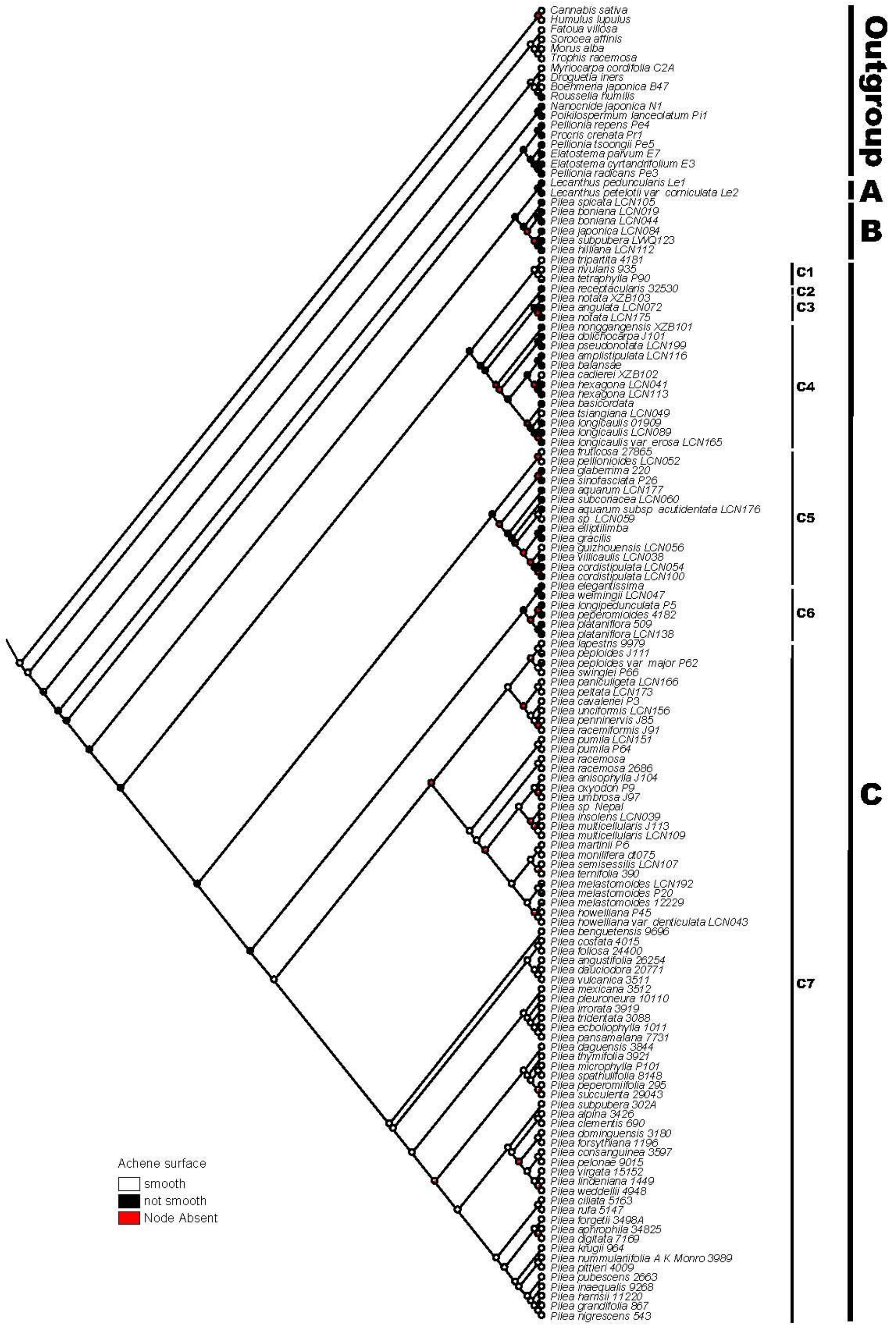


Fig. S21. Ancestral state reconstruction for *Pilea* based on Maximum likelihood analysis of achene surface.

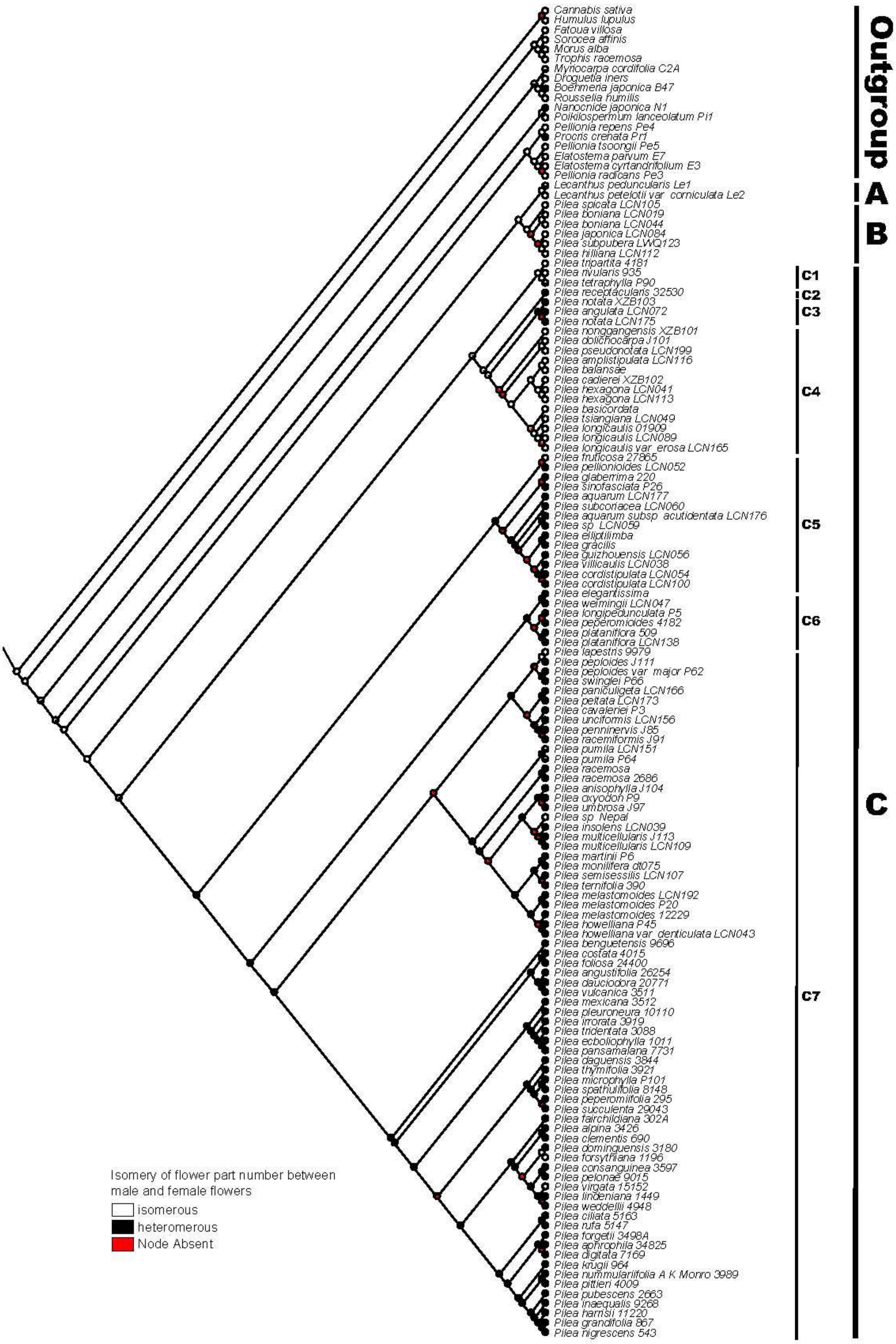


Fig. S22. Ancestral state reconstruction for *Pilea* based on Maximum likelihood analysis of isomery of flower part number between male and female flowers.

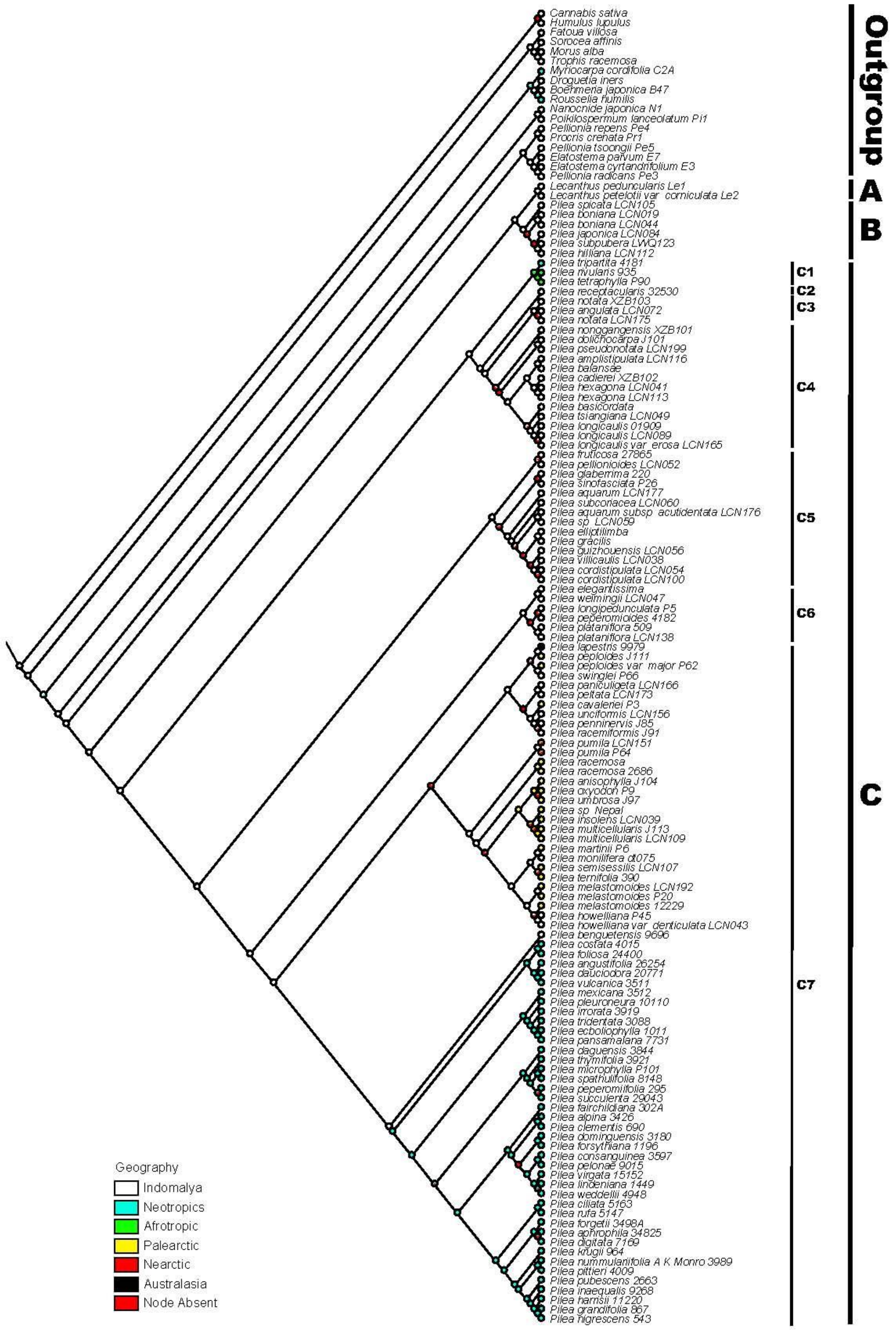


Fig. S23. Ancestral state reconstruction for *Pilea* based on Maximum likelihood analysis of geography.

#### Appendix A. Supplementary material

Supplementary Text 1. Species names and GenBank accession numbers of DNA sequences used in this study

Voucher information for samples of which sequences are newly generated is given using the following format: Taxon name, collection locality, collector and collector number (herbarium for voucher specimen), GenBank accession numbers for ITS, *trnL-trnF*, *rbcL*, respectively. Samples downloaded from NCBI only remain Taxon name, collection locality, GenBank accession numbers as stated above. Herbaria: IBK = Guangxi Institute of Botany; K = Royal Botanic Garden, Kew; KUN = Kunming Institute of Botany, Chinese Academy of Sciences; SING = Singapore Botanic Gardens. (NA = not available, * newly generated sequences).

Ingroup:

*Achudemia javanica*_LWQ123, Indonesia, Robinson H.C. and Kloss C.B. 1914 (SING), MT516339*, MT523094*, MT523050*. *Lecanthus peduncularis*_Le1, China, KF137871, KF138350, KF138186. *Lecanthus petelotii var corniculate*_Le2, China, KF137873, KF138352, KF138188. *Pilea alpina* 3426, Santo Domingo, DQ175543, DQ179309, NA. *Pilea amplistipulata*_LCN116, China, Shui Y.M. et. al.14319 (KUN), MT516340*, MT523095*, MT523051*. *Pilea angulata*_LCN072, China, Fu L.F. et al. FL0234 (IBK), MT516341*, MT523096*, MT523052*. *Pilea angustifolia*_26254, Costa Rica, DQ175556, DQ179289, NA. *Pilea anisophylla*_J104, China, Wen F. WF182817-10 (IBK), MT516342*, MT523097*, MT523053*. *Pilea aphrophila*_34825, Colombia, DQ175589, DQ179323, NA. *Pilea aquarum*_LCN177, China, Wei Y.G. Wei097 (IBK), MT516343*, MT523098*, MT523054*. *Pilea aquarum* subsp. *acutidentata*_LCN176, China, Wen F. WFLSH111207 (IBK), MT516344*, MT523099*, MT523055*. *Pilea balansae*, Vietnam, Huang S.L. HSL118-1 (IBK), MT516345*, MT523100*, NA. *Pilea basicordata*, China, DQ175614, DQ179361, NA. *Pilea benguetensis*_9696, Phillipines, DQ175554, DQ179337, NA. *Pilea boniana*_LCN019, China, Fu L.F. FLF180412-01, (IBK), MT516346*, MT523101*, MT523056*. *Pilea boniana*_LCN044, China, Qin et al. 3193 (KUN), MT516347*, MT523102*, MT523057*. *Pilea cadierei*_XZB102, China, Xin Z.B. XZB102 (IBK), MT516348*, MT523103*, MT523058*. *Pilea cavaleriei*_P3, China, KF137895, KF138380, KF138214. *Pilea ciliate*_5163, Jamaica, DQ175538, DQ179300, NA. *Pilea clementis*_690,

Cuba, DQ175550, DQ179310, NA. *Pilea consanguinea*_3597, Santo Domingo, DQ175539, DQ179312, NA. *Pilea cordistipulata*_LCN054, China, Monro et al. AM6727 (IBK), MT516349[*], MT523104[*], MT523059[*]. *Pilea cordistipulata*_LCN100, China, Huang S.L. HSL140 (IBK), MT516350[*], MT523105[*], MT523060[*]. *Pilea costata*_4015, Peru, DQ175595, DQ179290, NA. *Pilea daguensis*_3844, Mexico, DQ175567, DQ179332, NA. *Pilea dauciodora*_20771, Mexico, DQ175562, DQ176857, NA. *Pilea digitate*_7169, Panama, DQ175559, DQ179326, NA. *Pilea dolichocarpa*_J101, China, Monro A.K. AM6399 (IBK), MT516351[*], MT523106[*], MT523061[*]. *Pilea dominguensis*_3180, Santo Domingo, DQ175541, DQ179313, NA. *Pilea ecboliophylla*_1011, Mexico, DQ175531, DQ179292, NA. *Pilea elegantissima*_P39, China, MH357923, MH358303, MH358124. *Pilea elliptilimba*, China, Huang S.L. HSL113 (IBK), MT516352[*], MT523107[*], NA. *Pilea foliosa*_24400, Peru, DQ175571, DQ179291, NA. *Pilea forgetii*_3498A, Panama, DQ175585, DQ179333, NA. *Pilea forsythiana*_1196, Dominica, DQ175546, DQ179311, NA. *Pilea fruticose*_27865, Borneo, DQ175604, DQ179353, NA. *Pilea glaberrima*_220, Nepal, DQ175600, DQ179352, NA. *Pilea gracilis*, China, Wei Y.G. Wei039 (IBK), MT516353[*], MT523108[*], NA. *Pilea grandifolia* 867, Jamaica, DQ175551, DQ179303, NA. *Pilea guizhouensis*_LCN056, China, Monro et al. AM6715 (IBK), MT516354[*], MT523109[*], MT523062[*]. *Pilea harrisii* 11220, Jamaica, DQ175537, DQ179302, NA. *Pilea hexagona*_LCN041, China, Shui Y.M. YN004 (KUN), MT516355[*], MT523110[*], MT523063[*]. *Pilea hexagona*_LCN113, China, Sino-Vietnamese expidition 775 (KUN), MT516356[*], MT523111[*], MT523064[*]. *Pilea hilliana*_LCN112, China, liuzu 2014 (KUN), MT516357[*], MT523112[*], MT523065[*]. *Pilea howelliana* P45, China, MH357926, MH358306, MH358127. *Pilea howelliana var. denticulata*_LCN043, China, Wang Y.Z. 4678 (KUN), MT516358[*], MT523113[*], MT523066[*]. *Pilea inaequalis* 9268, Trinidad, DQ175552, DQ179304, NA. *Pilea insolens*_LCN039, China, FLPH Tibet Expedition 12-1838 (IBK), MT516359[*], MT523114[*], MT523067[*]. *Pilea irrorata* 3919, Mexico, DQ175535, DQ179294, NA. *Pilea japonica*_LCN084, China, Huang S.L. HSL012 (IBK), MT516360[*], MT523115[*], MT523068[*]. *Pilea krugii* 964, Puerto Rico, DQ175581, DQ179315, NA. *Pilea lapestris* 9979, Indonesia, DQ175598, DQ179341, NA. *Pilea lindeniana* 1449, Cuba, DQ175547, DQ179314, NA. *Pilea longicaulis* 01909, China, DQ175611, DQ179363, NA. *Pilea longicaulis var. erosa*_LCN165, China, Monro A.K.

AM6809 (IBK), MT516361[*], MT523116[*], NA. *Pilea longicaulis*_LCN089, China, Huang S.L. HSL120 (IBK), MT516362[*], MT523117[*], MT523069[*]. *Pilea longipedunculata*_P5, China, KF137897, KF138382, KF138216. *Pilea martini*_P6, China, KF137898, KF138383, KF138217. *Pilea melastomoides*_12229, Indonesia, DQ175596, DQ179345, NA. *Pilea melastomoides*_P20, China, KF137899, KF138384, KF138218. *Pilea melastomoides*_LCN192, China, Huang S.L. HSL124 (IBK), MT516363[*], MT523118[*], MT523070[*]. *Pilea Mexicana*_3512, Panama, DQ175579, DQ179278, NA. *Pilea microphylla*_P101, Brazil, MH357928, MH358307, MH358129. *Pilea monilifera*_dt075, China, MK911055, MK911077, MK911100. *Pilea multicellularis*_J113, China, Wen F. WF180817-08 (IBK), MT516364[*], MT523119[*], MT523071[*]. *Pilea multicellularis*_LCN109, China, Tibet Expedition 12-1247 (IBK), MT516365[*], MT523120[*], MT523072[*]. *Pilea nigrescens*_543, Jamaica, DQ175582, DQ179301, NA. *Pilea nonggangensis*_XZB101, China, Huang S.L. HSL149 (IBK), MT516366[*], MT523121[*], MT523073[*]. *Pilea notata*_LCN175, China, Wen F. WFLSH120925 (IBK), MT516367[*], MT523122[*], MT523074[*]. *Pilea notata*_XZB103, China, Huang S.L. HSL099 (IBK), MT516368[*], MT523123[*], MT523075[*]. *Pilea nummularifolia* A.K.Monro 3989, Peru, DQ175588., DQ179316, NA. *Pilea oxyodon* P9, China, KF137902, KF138387, KF138221. *Pilea paniculigera*_LCN166, China, Monro A.K. AM6818 (IBK), MT516369[*], MT523124[*], MT523076[*]. *Pilea pansamalana* 7731, Mexico, DQ175533, DQ179296, NA. *Pilea pellionioides*_LCN052, China, Hu G.X. HGX001 (IBK), MT516370[*], MT523125[*], MT523077[*]. *Pilea pelonae* 9015, Dominican, DQ175540, DQ179327, NA. *Pilea peltata*_LCN173, China, Huang S.L. HSL116 (IBK), MT516371[*], MT523126[*], MT523078[*]. *Pilea penninervis*_J85, China, Wei Y.G. 0726 (IBK), MT516372[*], MT523127[*], MT523079[*]. *Pilea peperomiifolia*_295, Virgin Islands, DQ175569, DQ179281, NA. *Pilea peperomioides*_4182, cultivated in UK, DQ175605, DQ179350, NA. *Pilea peploides var. major*_P62, China, MH357931, NA, MH358132. *Pilea peploides*_J111, China, Monro A.K. AM6433 (IBK), MT516373[*], MT523128[*], MT523080[*]. *Pilea pittieri*_4009, Peru, DQ175560, DQ179328, NA. *Pilea plataniflora*_509, Japan, DQ175599, DQ179349, NA. *Pilea plataniflora*_LCN138, China, Huang S.L. HSL018 (IBK), MT516374[*], MT523129[*], MT523081[*]. *Pilea pleuroneura*_10110, Guatemala, DQ175532, DQ179297, NA. *Pilea pseudonotata*_LCN199, China, Wei Y.G. Wei047 (IBK), MT516375[*], MT523130[*],

MT523082[*]. *Pilea pubescens*_2663, Belize, DQ175558, DQ179325, NA. *Pilea pumila*_P64, China, MH357932, MH358309, MH358133. *Pilea pumila*_LCN151, China, Huang S.L. 2104H (IBK), MT516376[*], MT523131[*], MT523083[*]. *Pilea racemiformis*_J91, China, Wen F. WF150423-33 (IBK), MT516377[*], MT523132[*], MT523084[*]. *Pilea racemosa*, China, DQ175602, DQ179347, NA. *Pilea racemose*_2686, China, DQ175602, DQ179347, NA. *Pilea receptacularis*_32530, China, DQ175612, DQ179362, NA. *Pilea rivularis*_935, Tanzania, DQ175606, DQ179358, NA. *Pilea rufa*_5147, Jamaica, DQ175578, DQ179299, NA. *Pilea semisessilis*_LCN107, China, Zhou Z.K. et al. EXLS-0272 (KUN), MT516378[*], MT523133[*], MT523085[*]. *Pilea sinofasciata*_P26, China, KF137905, KF138389, KF138224. *Pilea* sp. _LCN059, China, Fu L.F. FLF180527-01 (IBK), MT516379[*], MT523134[*], MT523086[*]. *Pilea* sp. _Nepal, cultivated at RBGE, DQ175601, DQ179344, NA. *Pilea spathulifolia*_8148, Dominican Republic, DQ175570, DQ179282, NA. *Pilea spicata*_LCN105, China, Burkill H. 37636, (K), MT516380[*], MT523135[*], MT523087[*]. *Pilea subcoriacea*_LCN060, China, Wei Y.G. WYG180519-02 (IBK), MT516381[*], MT523136[*], MT523088[*]. *Pilea succulenta*_29043, Cayman Islands, DQ175565, DQ179280, NA. *Pilea swinglei*_P66, China, MH357933, NA, MH358134. *Pilea ternifolia*_390, Nepal, DQ175597, DQ179346, NA. *Pilea tetraphylla*_P90, Madagascar, MH357934, MH358310, MH358135. *Pilea thymifolia*_3921, Peru, DQ175568, DQ179283, NA. *Pilea tridentata*_3088, Mexico, DQ175536, DQ179293, NA. *Pilea tripartita*_4181, Panama, DQ175617, DQ176859, NA. *Pilea tsiangiana*_LCN049, China, Monro A.K. AM6770 (IBK), MT516382[*], MT523137[*], MT523089[*]. *Pilea umbrosa*_J97, China, Wen F. WF180821-01 (IBK), MT516383[*], MT523138[*], MT523090[*]. *Pilea unciformis*_LCN156, China, Huang S.L. HSL132 (IBK), MT516384[*], MT523139[*], MT523091[*]. *Pilea villicaulis*_LCN038, China, Shui et al. 12871 (KUN), MT516385[*], MT523140[*], MT523092[*]. *Pilea virgata*_15152, Jamaica, DQ175548, DQ179329, NA. *Pilea vulcanica*_3511, Panama, DQ175563, DQ179284, NA. *Pilea weddellii*_4948, Jamaica, DQ175545, DQ179308, NA. *Pilea weimingii*_LCN047, China, Lv R.D. LRD001 (IBK), MT516386[*], MT523141[*], MT523093[*]. *Pilea fairchildiana*_302A, Dominica, JN252482, JN252481, NA.

Outgroup:

*Boehmeria japonica*_B47, USA, KF137808, KF138279, KF138116. *Cannabis sativa*_Z1773,

China, MH357863, NA, MH358052. *Droguetia iners*_Dr1, China, KF137844, KF138318, KF138154. *Elatostema cyrtandrifolium*_E3, China, KF137848, KF138322, KF138158. *Elatostema parvum*_E7, China, KF137852, KF138326, KF138162. *Fatoua villosa*_F1, China, KF137858, KF138331, KF138168 . *Humulus lupulus*_D3848, China, MH357893, NA, MH358086. *Morus alba*, HM747164, HM747180, L01933. *Myriocarpa cordifolia*_C2A, Panama, KF137877, KF138357, KF138193. *Nanocnide japonica*_N1, China, KF137879, KF138359, KF138194. *Pellionia radicans*_Pe3, China, KF137891, KF138375, KF138210. *Pellionia repens*_Pe4, China, KF137892, KF138376, KF138211. *Pellionia tsoongii*_Pe5, China, KF137893, KF138377, KF138212. *Procris crenata*_Pr1, China, KF137922, KF138407, KF138242. *Sorocea affinis*, HM747179, HM747195, GQ981880. *Trophis racemosa*, HM747178, HM747194, GQ981908. *Poikilospermum lanceolatum*_Pi1, China, KF137912, KF138396, KF138231. *Rousselia humilis*, Dominica, KM586474, KM586646, KM586560.

### Supplementary Text 2. Morphological trait matrix.

| Spcecies\trait | Habit<br>terrestrial (0)<br>epiphytic or epipetric (1) | Plants lignified<br>lignified (0)<br>not lignified (1) | Plants with a dominant axis<br>with a single dominant axis (0)<br>lacking a single dominant axis (1) | Leaves arrangement<br>borne in the same plane (0)<br>borne spirally (1) | Leaf margin (excluding apex)<br>entire (0)<br>not entire (1) | Leaf venation<br>basal pair of lateral veins not substantially thicker than subsequent pairs (pinnately veined) (0)<br>basal pair of lateral veins substantially thicker than subsequent pairs (3-veined) (1) | Stipule length<br>≤10 (0)<br>>10 (1) | Staminate inflorescence architecture<br>branched (0)<br>not branched (1) | Staminate branched inflorescence type<br>branches not strongly reduced forming a capitate head (0)<br>branches strongly reduced forming a capitate or fascicle-like head (1)<br>spicate / racemose (NA) | Staminate inflorescence symmetry<br>symmetrical (0)<br>asymmetrical (1) | Receptacle in staminate inflorescence<br>basal branches of inflorescence not fused to form a receptacle-like structure (0)<br>some basal branches of inflorescence fused to form a receptacle-like structure (1) | Staminate sepal number<br>2 (0)<br>3 (1)<br>4 (2)<br>5 (3) |
|---|---|---|---|---|---|---|---|---|---|---|---|---|
| Boehmeria japonica B47 | 0 | 0/1 | 1 | 0 | 1 | 1 | 0 | 0 | 0 | 1 | 0 | 2 |
| Cannabis sativa | NA | NA | NA | NA | NA | NA | NA | NA | NA | NA | NA | NA |
| Droguetia iners | 0 | 1 | 1 | 0 | 1 | 1 | 0 | 1 | NA | 0 | 1 | 0 |
| Elatostema cyrtandrifolium E3 | 0/1 | 1 | 1 | 0 | 1 | 1 | 0 | 1 | NA | 0 | 1 | 2 |
| Elatostema parvum E7 | 0 | 1 | 1 | 0 | 1 | 1 | 0 | 1 | NA | 0 | 1 | 3 |
| Fatoua villosa | NA | NA | NA | NA | NA | NA | NA | NA | NA | NA | NA | NA |
| Humulus lupulus | NA | NA | NA | NA | NA | NA | NA | NA | NA | NA | NA | NA |
| Lecanthus peduncularis Le1 | 0/1 | 1 | 1 | 0 | 1 | 1 | 0 | 1 | NA | 0 | 1 | 3 |
| anthus petelotii var corniculata | 0/1 | 1 | 1 | 0 | 1 | 1 | 0 | 1 | NA | 0 | 1 | 2 |
| Morus alba | NA | NA | NA | NA | NA | NA | NA | NA | NA | NA | NA | NA |
| Myriocarpa_cordifolia_C2A | 0 | 0 | 0 | 1 | 1 | 1 | 1 | 1 | NA | 0 | 0 | 2 |
| Nanocnide japonica N1 | 0 | 1 | 1 | 0 | 1 | 1 | 0 | 0 | 0 | 1 | 0 | 3 |
| Pellionia radicans Pe3 | 0/1 | 1 | 1 | 0 | 1 | 1 | 0 | 0 | 0 | 0 | 0 | 3 |
| Pellionia repens Pe4 | 0/1 | 1 | 1 | 0 | 1 | 1 | 1 | 0 | 0 | 1 | 0 | 3 |
| Pellionia tsoongii Pe5 | 0/1 | 0/1 | 1 | 0 | 0 | 0 | 1 | 0 | 0 | 1 | 0 | 3 |
| Pilea alpina 3426 | 0 | 0 | 1 | 0 | 0 | 1 | 0 | 0 | 1 | 0 | 0 | 2 |
| Pilea amplistipulata_LCN116 | 1 | 0 | 1 | 0 | 0/1 | 1 | 1 | 0 | 0 | 1 | 0 | 2 |
| Pilea angulata_LCN072 | 0 | 1 | 1 | 0 | 1 | 1 | 1 | 0 | 0 | 1 | 0 | 2 |
| Pilea angustifolia 26254 | 0 | 0 | 0 | 0 | 1 | 1 | 0/1 | 0 | 1 | 0 | 0 | 2 |
| Pilea anisophylla_J104 | 0 | 1 | 1 | 0 | 0/1 | 1 | 0 | 1 | NA | 0 | 0 | 2 |
| Pilea aphrophia 34825 | 1 | 1 | 1 | 0 | 1 | 1 | 0 | 0 | 1 | 0 | 0 | 2 |
| Pilea aquarum LCN177 | 0 | 1 | 1 | 0 | 1 | 1 | 0 | 0 | 0 | 1 | 0 | 2 |
| quarum subsp. acutidentata_L | 0 | 1 | 1 | 0 | 1 | 1 | 0 | 0 | 0 | 1 | 0 | 2 |
| Pilea balansae | 1 | 0 | 0 | 0 | 0/1 | 1 | 1 | 0 | 0 | 1 | 0 | 2 |
| Pilea basicordata | 1 | 0 | 0 | 0 | 1 | 1 | 1 | 0 | 0 | 1 | 0 | 2 |
| Pilea benguetensis 9696 | 0 | 0 | 0 | 0 | 1 | 1 | 0 | 0 | 1 | 1 | 0 | 2 |
| Pilea boniana_LCN019 | 1 | 1 | 1 | 0 | 1 | 1 | 1 | 0 | 0 | 1 | 0 | 3 |
| Pilea boniana_LCN044 | 1 | 1 | 1 | 0 | 1 | 1 | 1 | 0 | 0 | 1 | 0 | 3 |
| Pilea cadierei_XZB102 | 0 | 0/1 | 1 | 0 | 1 | 1 | 1 | 0 | 1 | 0 | 0 | 2 |
| Pilea cavaleriei P3 | 1 | 1 | 1 | 0 | 0/1 | 1 | 0 | 0 | 1 | 0 | 0 | 2 |
| Pilea ciliata 5163 | 0 | 0 | 1 | 0 | 1 | 1 | 0 | 0 | 1 | 0 | 0 | 2 |
| Pilea clementis 690 | 0 | 0 | 1 | 0 | 0 | 1 | 0 | 0 | 1 | 0 | 0 | 2 |
| Pilea consanguinea 3597 | 0 | 0 | 0 | 0 | 0 | 1 | 0 | NA | NA | NA | NA | NA |
| Pilea cordistipulata_LCN054 | 0 | 1 | 1 | 0 | 1 | 1 | 0 | 0 | 0 | 1 | 0 | 2 |
| Pilea cordistipulata_LCN100 | 0 | 1 | 1 | 0 | 1 | 1 | 0 | 0 | 0 | 1 | 0 | 2 |
| Pilea costata 4015 | 1 | 0 | 0 | 0 | 1 | 1 | 0 | NA | NA | NA | NA | NA |
| Pilea daguensis 3844 | 0/1 | 1 | 0 | 0 | 1 | 1 | 0 | 0 | 1 | 0 | 0 | 2 |
| Pilea dauciodora 20771 | 0 | 1 | 1 | 0 | 1 | 1 | 0/1 | 0 | 1 | 1 | 0 | 2 |
| Pilea digitata 7169 | 1 | 0 | 0 | 0 | 1 | 1 | 0 | 0 | 1 | 0 | 0 | 2 |
| Pilea dolichocarpa_J101 | 0 | 0/1 | 1 | 0 | 1 | 1 | 0 | 0 | 0 | 0 | 0 | 2 |
| Pilea dominguensis 3180 | 0 | 0 | 0 | 0 | 1 | 1 | 0 | 0 | 1 | 1 | 0 | 2 |
| Pilea ecboliophylla 1011 | 1 | 0 | 1 | 0 | 1 | 1 | 0 | 0 | 0 | 0 | 0 | 2 |
| Pilea elegantissima | 1 | 1 | 1 | 0 | 0 | 1 | 0 | 1 | NA | 0 | 0 | 2 |
| Pilea elliptilimba | 0 | 1 | 1 | 0 | 1 | 1 | 0 | NA | NA | NA | NA | NA |
| Pilea fairchildiana_302A | 0 | 0 | 1 | 1 | 0 | NA (no lateral nerves) | 1 | 0 | 0 | 1 | 0 | 2 |
| Pilea foliosa 24400 | 1 | 0 | 1 | 0 | 0 | 0 | 0 | NA | NA | NA | NA | 2 |
| Pilea forgetii 3498A | 1 | 0 | 0 | 0 | 1 | 1 | 1 | 0 | 1 | 0 | 0 | 2 |
| Pilea forsythiana 1196 | 0 | 0 | 1 | 0 | 0 | 1 | 0 | NA | NA | NA | NA | NA |
| Pilea fruticosa 27865 | 0 | 0 | 1 | 0 | 1 | 1 | 0 | 0 | 0 | 1 | 0 | NA |
| Pilea glaberrima 220 | 0 | 0/1 | 1 | 0 | 1 | 1 | 0 | 0 | 0 | 1 | 0 | 2 |
| Pilea gracilis | 0 | 1 | 1 | 0 | 1 | 1 | 0 | 0 | 0 | 1 | 0 | 2 |
| Pilea grandiflolia 867 | 0 | 0 | 0 | 0 | 1 | 1 | 1 | 0 | 0 | 1 | 0 | 2 |
| Pilea guizhouensis_LCN056 | 0/1 | 0/1 | 1 | 0 | 1 | 1 | 0 | 0 | 0 | 0 | 0 | 2 |
| Pilea harrisii 11220 | 0/1 | 1 | 1 | 0 | 1 | 1 | 0 | 0 | 1 | 0 | 0 | 2 |
| Pilea hexagona_LCN041 | 1 | 0 | 1 | 0 | 1 | 1 | 1 | 0 | 1 | 0 | 0 | 2 |
| Pilea hexagona_LCN113 | 1 | 0 | 1 | 0 | 1 | 1 | 1 | 0 | 1 | 0 | 0 | 2 |
| Pilea hilliana_LCN112 | 0 | 1 | 1 | 0 | 1 | 1 | 0 | 0 | 0 | 0 | 0 | 3 |
| Pilea howelliana P45 | 0 | 1 | 1 | 0 | 1 | 1 | 0 | 0 | 0 | 1 | 0 | 2 |
| howelliana var. denticulata_LC | 0 | 1 | 1 | 0 | 1 | 1 | 0 | 0 | 0 | 1 | 0 | 2 |
| Pilea inaequalis 9268 | 0 | 1 | 0 | 0 | 1 | 1 | 0 | 0 | 0 | 1 | 0 | 2 |
| Pilea insolens_LCN039 | 0 | 1 | 1 | 0 | 0 | 1 | 0 | 1 | NA | 0 | 0 | 2 |
| Pilea irrorata 3919 | 0 | 0 | 0 | 0 | 1 | 1 | 0 | 0 | 1 | 0 | 0 | 2 |
| Pilea japonica_LCN084 | 1 | 1 | 1 | 0 | 1 | 1 | 1 | 0 | 1 | 1 | 0 | 3 |
| Pilea krugii 964 | 1 | 0 | 1 | 0 | 1 | 0 | 0 | 0 | 1 | 1 | 0 | 2 |
| Pilea lapestris 9979 | 0 | 0 | 1 | 0 | 0 | NA | 0 | 1 | NA | NA | NA | 2 |
| Pilea lindeniana 1449 | 0 | 0 | 0 | 0 | 0 | 1 | 0 | 0 | 0 | 1 | 0 | 2 |
| Pilea longicaulis 01909 | 0 | 0 | 1 | 0 | 0/1 | 1 | 0 | 0 | 0 | 1 | 0 | 2 |
| ea longicaulis var. erosa_LCN1 | 0 | 0 | 1 | 0 | 1 | 1 | 0 | 0 | 0 | 1 | 0 | 2 |
| Pilea longicaulis_LCN089 | 0 | 0 | 1 | 0 | 0/1 | 1 | 0 | 0 | 0 | 1 | 0 | 2 |
| Pilea longipedunculata P5 | 0 | 1 | 1 | 0 | 0 | 1 | 0 | 0 | 0 | 1 | 0 | 2 |
| Pilea martinii P6 | 0 | 1 | 1 | 0 | 1 | 1 | 0 | 0 | 0 | 1 | 0 | 2 |
| Pilea melastomoides 12229 | 0 | 1 | 1 | 0 | 1 | 1 | 0 | 0 | 0 | 1 | 0 | 2 |
| Pilea melastomoides P20 | 0 | 1 | 1 | 0 | 1 | 1 | 0 | 0 | 0 | 1 | 0 | 2 |
| Pilea melastomoides_LCN192 | 0 | 1 | 1 | 0 | 1 | 1 | 0 | 0 | 0 | 1 | 0 | 2 |
| Pilea mexicana 3512 | 0 | 0 | 0 | 0 | 0 | NA (no lateral nerves) | 0 | 0 | 1 | 0 | 0 | 2 |
| Pilea microphylla P101 | 0/1 | 1 | 1 | 0 | 0 | 0 | 0 | 0 | 1 | 0 | 0 | 2 |
| Pilea monilifera dt075 | 0 | 1 | 1 | 0 | 1 | 1 | 0 | 1 | NA | 0 | 0 | 2 |
| Pilea multicellularis_J113 | 0 | 1 | 1 | 0 | 1 | 1 | 0 | 0 | 0 | 1 | 0 | 2 |
| Pilea multicellularis_LCN109 | 0 | 1 | 1 | 0 | 1 | 1 | 0 | 0 | 0 | 1 | 0 | 2 |
| Pilea nigrescens 543 | 0 | 0 | 1 | 0 | 1 | 1 | 0 | 0 | 1 | 0 | 0 | 2 |
| Pilea nonggangensis_XZB101 | 1 | 0 | 0 | 0 | 0/1 | 1 | 1 | 0 | 0 | 1 | 0 | 2 |
| Pilea notata_LCN175 | 0 | 1 | 1 | 0 | 1 | 1 | 1 | 0 | 0 | 1 | 0 | 2 |
| Pilea notata_XZB103 | 0 | 1 | 1 | 0 | 1 | 1 | 1 | 0 | 0 | 1 | 0 | 2 |
| a nummulariifolia A.K.Monro | 0/1 | 1 | 1 | 1 | 1 | 1 | 0 | 0 | 0/1 | 0 | 0 | 2 |
| Pilea oxyodon P9 | 0 | 1 | 1 | 0 | 1 | 1 | 0 | 0 | 0 | 1 | 0 | 2 |
| Pilea paniculigera_LCN166 | 1 | 1 | 1 | 0 | 1 | 1 | 0 | 0 | 0 | 1 | 0 | 2 |
| Pilea pansamalana 7731 | 1 | 0 | 1 | 0 | 1 | 1 | 0 | 0 | 0 | 0 | 0 | 2 |
| Pilea pellionioides_LCN052 | 0 | 0/1 | 1 | 0 | 1 | 1 | 0 | 0 | 0 | 1 | 0 | 2 |
| Pilea pelonae 9015 | 1 | 0 | 1 | 0 | 0 | NA (no lateral nerves) | 0 | 0 | 1 | 0 | 0 | 2 |
| Pilea peltata_LCN173 | 1 | 1 | 1 | 0 | 1 | 1 | 0 | 1 | NA | 0 | 0 | 2 |
| Pilea penninervis_J85 | 0 | 1 | 1 | 0 | 1 | 0 | 0 | 1 | NA | 0 | 0 | 2 |
| Pilea peperomiifolia 295 | 1 | 1 | 1 | 0 | 0 | NA (no lateral nerves) | 0 | 0 | 1 | 0 | 0 | 2 |
| Pilea peperomioides 4182 | 0 | 1 | 1 | 1 | 0 | 1 | 0/1 | 0 | 0 | 1 | 0 | 2 |
| Pilea peploides var. major 62 | 0 | 1 | 1 | 0 | 1 | NA | 0 | 0 | 1 | 0 | 0 | 2 |
| Pilea peploides_J111 | 0 | 1 | 1 | 0 | 1 | NA | 0 | 0 | 1 | 0 | 0 | 2 |
| Pilea pittieri 4009 | 0/1 | 0 | 0 | 0 | 1 | 1 | 0/1 | 0 | 1 | 1 | 0 | 2 |
| Pilea plataniflora 509 | 1 | 1 | 1 | 0 | 0 | 1 | 0 | 0 | 0 | 1 | 0 | 2 |
| Pilea plataniflora_LCN138 | 1 | 1 | 1 | 0 | 0 | 1 | 0 | 0 | 0 | 1 | 0 | 2 |
| Pilea pleuroneura 10110 | 0 | 0 | 0 | 0 | 1 | 0 | 0 | 0 | 1 | 0 | 0 | 2 |
| Pilea pseudonotata_LCN199 | 0 | 0 | 1 | 0 | 1 | 1 | 1 | 0 | 0 | 1 | 0 | 2 |
| Pilea pubescens 2663 | 0 | 1 | 0 | 0 | 1 | 1 | 0 | 0 | 0 | 1 | 0 | 2 |
| Pilea pumila P64 | 0 | 1 | 1 | 0 | 1 | 1 | 0 | 0 | 0 | 1 | 0 | 0/1/2 |
| Pilea pumila_LCN151 | 0 | 1 | 1 | 0 | 1 | 1 | 0 | 0 | 0 | 1 | 0 | 0/1/2 |
| Pilea racemiformis_J91 | 1 | 1 | 1 | 0 | 1 | 1 | 0 | 0 | 0 | 1 | 0 | 2 |
| Pilea racemosa | 0 | 1 | 1 | 0 | 1 | 1 | 0 | 0/1 | 0/1 | 0/1 | 0 | 2 |
| Pilea racemosa 2686 | 0 | 1 | 1 | 0 | 1 | 1 | 0 | 0/1 | 0/1 | 0/1 | 0 | 2 |
| Pilea receptacularis 32530 | 0 | 1 | 1 | 0 | 1 | 1 | 0 | 1 | NA | 0 | 1 | 2 |
| Pilea rivularis 935 | 0 | 1 | 1 | 0 | 1 | 1 | 0 | 0 | 1 | 0 | 1 | 1 |
| Pilea rufa 5147 | 0 | 0 | 0 | 0 | 1 | 1 | 0 | 0 | 1 | 0 | 0 | 2 |
| Pilea semisessilis_LCN107 | 0 | 1 | 1 | 0 | 1 | 1 | 0 | 0 | 0 | 1 | 0 | 2 |
| Pilea sinofasciata P26 | 0 | 1 | 1 | 0 | 1 | 1 | 0 | 0 | 0 | 1 | 0 | 2 |
| Pilea sp. LCN059 | 0/1 | 1 | 1 | 0 | 1 | 1 | 0 | NA | NA | NA | NA | NA |
| Pilea sp. Nepal | NA | NA | NA | NA | NA | NA | NA | NA | NA | NA | NA | NA |
| Pilea spathulifolia 8148 | 0 | 0 | 0 | 0 | 0 | NA (no lateral nerves) | 0 | 0 | 1 | 0 | 0 | 2 |
| Pilea spicata_LCN105 | 0 | 1 | 1 | 0 | 1 | 1 | 1 | NA | NA | NA | NA | NA |
| Pilea subcoriacea_LCN060 | 0 | 1 | 1 | 0 | 1 | 1 | 0 | 0 | 0 | 1 | 0 | 2 |
| Pilea subpubera_LWQ123 | NA | 1 | 1 | 0 | 1 | 1 | 0 | 0 | 0 | 1 | 0 | 3 |
| Pilea succulenta 29043 | 0 | 0 | 0 | 0 | 1 | 1 | 1 | NA | NA | NA | NA | NA |
| Pilea swinglei P66 | 0 | 1 | 1 | 0 | 1 | 1 | 0 | 0/1 | 0 | 0/1 | 0 | 2 |
| Pilea ternifolia 390 | 0 | 1 | 1 | 0 | 1 | 0 | 0 | 0 | 1 | 0 | 0 | 2 |
| Pilea tetraphylla P90 | 0 | 1 | 1 | 0 | 1 | 1 | 0 | 0 | 0 | 1 | 0 | 0 |
| Pilea thymifolia 3921 | 0/1 | 0 | 0 | 0 | 0 | NA (no lateral nerves) | 0 | NA | 1 | 0 | 0 | 2 |
| Pilea tridentata 3088 | 0/1 | 0 | 0 | 0 | 0 | 0 | 0 | 0 | 0 | 0 | 0 | 2 |
| Pilea tripartita 4181 | 0 | 1 | 1 | 0 | 1 | 1 | 0 | 0 | NA | 0 | 1 | 1 |
| Pilea tsiangiana_LCN049 | 0 | 0 | 1 | 0 | 1 | 1 | 0 | 0 | 0 | 1 | 0 | 2 |
| Pilea umbrosa J97 | 0 | 1 | 1 | 0 | 1 | 1 | 0 | 0 | 0 | 1 | 0 | 2 |
| Pilea unciformis_LCN156 | 1 | 1 | 1 | 0 | 1 | 1 | 0 | 0 | 0 | 0 | 0 | 2 |
| Pilea villicaulis_LCN038 | 0 | 1 | 1 | 0 | 1 | 1 | 0 | 0 | 0 | 1 | 0 | 2 |
